# Triple-Negative Breast Cancer Cells Activate Sensory Neurons via TRPV1 to Drive Nerve Outgrowth and Tumor Progression

**DOI:** 10.1101/2025.07.22.666151

**Authors:** Hanan Bloomer, Savannah R. Parker, Audrey L. Pierce, Wesley Clawson, Mallory M. Caron, Thomas Gerton, Ankit Pandey, Thanh T. Le, Tolulope Adewumi, Charlotte Kuperwasser, Michael Levin, Madeleine J. Oudin

**Affiliations:** Department of Developmental, Molecular & Chemical Biology, Tufts University School of Medicine, Boston, MA 02111, USA; Department of Biomedical Engineering, Tufts University, Medford, MA 02155, USA; Department of Biology, Tufts University, Medford, MA 02155, USA; Vinmec-VinUni Institute of Immunology, College of Health Science, VinUniversity, Hanoi, Vietnam; Laboratory for the Convergence of Biomedical, Physical, and Engineering Sciences, Tufts University School of Medicine, Boston, MA 02111, USA; Allen Discovery Center, Tufts University, Medford, MA 02155, USA; Wyss Institute for Biologically Inspired Engineering, Harvard University, Boston, MA 02115, USA

## Abstract

The tumor microenvironment in triple-negative breast cancer (TNBC) is characterized by increased sensory nerve density, which contributes to cancer progression by promoting migration and metastasis. However, the origin of tumor-innervating nerves and the mechanisms driving sensory innervation into tumors remain poorly understood. Using *in vivo* retrograde labeling techniques, we show that mammary tumors are associated with an increase in nerves originating from the dorsal root ganglia. Additionally, we find that TNBC cells trigger stress and activation markers and induce neuronal firing in a transient receptor potential anilloid subtype 1 (TRPV1)-dependent manner. In both 2D and 3D cell culture models, TNBC cells promote outgrowth of sensory nerves that is abrogated with *Trpv1* knockout. We identified c-Jun and IL-6 as an effector of neurite outgrowth that acts downstream of TRPV1 to promote outgrowth *in vitro*. Finally, in *Trpv1* knockout mice, TNBC tumors exhibit delayed growth and reduced lung metastasis. These findings suggest that a critical role for TRPV1 in tumor-nerve crosstalk, offering a potential target to reduce metastatic disease.

## Introduction

The tumor microenvironment (TME) plays a critical role in driving cancer progression by influencing cell growth and metastasis. The complex and continuously evolving TME is composed of the extracellular matrix (ECM), a heterogenous collection of resident and infiltrating cells, and blood and lymphatic vessels^1^. Nerves were first documented to infiltrate the TME more than 100 years ago^2^. Since then, increased peripheral nerve density has been identified in a host of solid tumors, including breast, prostate, colorectal, lung, pancreatic, ovarian, and head and neck carcinomas^3–9^. The peripheral nervous system is comprised of sympathetic, parasympathetic, sensory, and motor nerves. Presence of all but the latter have been identified in tumors, with some tumors having a single innervation source, while others are composed of several nerve types^10, 11^. More recently, nerves have been shown to regulate tumor progression, including both promoting and suppressing tumor growth and metastasis. Understanding the mechanisms driving increased tumor innervation could provide a novel strategy to reduce metastatic disease and improve patient survival.

Breast cancer is the most common form of malignancy in women, accounting for almost one in three new female cancer diagnoses each year^12^. Metastatic disease, where tumor cells disseminate to secondary sites in the body, remains the primary cause of breast cancer-related deaths, resulting in 44,000 deaths annually in the United States^13^. Triple-negative breast cancer (TNBC) is a highly invasive molecular subtype characterized by the lack of targetable cell surface receptors and high rates of distant organ metastasis^14^. Human breast tumors have an increase in sensory nerve fiber density compared to healthy breast tissue, and TNBC exhibits increased innervation compared to other subtypes^5, 15, 16^. Across all subtypes, innervation is associated with higher rates of distant organ metastasis and decreased disease-free survival^5, 17, 18^. Co-injection of TNBC cells and sensory neurons into the mammary fat pads of mice, which leads to increased sensory nerve density in the primary tumor, significantly increases tumor growth and metastasis, while chemical denervation of sensory nerves reduces tumor progression^15, 16^.

Sensory nerves retain the ability to grow and regenerate into adulthood, suggesting that increased sensory nerve innervation in TNBC is driven by the outgrowth of existing nerves from the healthy adjacent tissue into the tumor. This growth is regulated by extrinsic cues, including neurotrophic factors (e.g., NGF, BDNF), axon guidance molecules (e.g., semaphorins, netrins), ECM components, and bioelectric signals such as spontaneous or evoked neural activity^19, 20^. These cues act through receptors like Trk receptors and ion channels, triggering intracellular pathways that activate transcriptional programs driving cytoskeletal remodeling, membrane trafficking, and neurite extension. While these pathways have been well characterized in development and injury, their role in tumor-induced nerve growth is only beginning to be explored. Deletion of the axon guidance molecule SLIT2 in endothelial cells significantly reduced sensory nerve density in a 4T1 xenograft mouse model of TNBC^16^. In rats, pharmacologic inhibition of VEGF or nerve growth factor (NGF) receptors in a N-Methyl-N-nitrosourea (MNU)-induced model of breast cancer decreased tumoral sensory nerve innervation^21^. However, the direct effects of tumor cells themselves on sensory nerves and the downstream neuronal mechanisms mediating tumor innervation remain poorly understood.

Transient receptor potential vanilloid subtype 1 (TRPV1) is a non-selective cation channel predominantly expressed on peripheral sensory neurons that plays a key role in detecting noxious stimuli^22^. Its activation leads to the influx of calcium and sodium ions, resulting in membrane depolarization and the transmission of pain signals to the central nervous system. In addition to its role in nociception, TRPV1 contributes to neurite outgrowth and regeneration through calcium-dependent signaling pathways^23–25^. Many known activators and sensitizers of TRPV1, such as heat (>43°C), mild acidification, inflammatory mediators, and NGF, are secreted by TNBC cells or present in the TME^5, 26, 27^. High TRPV1 expression in TNBC is associated with decreased patient survival^15^. Based on these observations, we hypothesized that tumor cell-derived secreted factors activate TRPV1 on sensory nerves in adjacent healthy tissue, triggering regenerative signaling that promotes neurite outgrowth into the TME.

Here, we show that mammary tumors display increased innervation by nerves originating from the dorsal root ganglia along the spinal cord. Using methods novel to the cancer neuroscience field, including microelectrode arrays and 3D hydrogel models, we demonstrate that TNBC cells stimulate sensory nerve activation and neurite outgrowth through TRPV1 signaling. Additionally, we find that cJun-mediated IL-6 expression acts downstream of TRPV1 to promote neurite outgrowth. Finally, genetic knockout of *Trpv1 in vivo* delays tumor growth and reduces lung metastasis. Together, our results provide novel insight into how tumors recruit nerves, which then drive increased tumor growth and progression.

## Results

### Tumor innervating nerves originate in the dorsal root ganglia

Increased sensory nerve density in TNBC is likely driven by the outgrowth of existing nerves into the tumor^10^. As healthy breast tissue is primarily innervated by sensory neurons originating from the dorsal root ganglia (DRG), we hypothesized that tumor-innervating nerves originate from pre-existing DRG neurons that extend into the TME^28^. Previous studies have demonstrated the presence of DRG nerves in both mouse and rat models of breast cancer using retrograde labeling techniques^16, 21^. However, these studies lacked non-tumor controls, making it difficult to assess how sensory innervation from DRG neurons evolves during tumor progression relative to normal mammary tissue. To address this, we injected the retrograde tracer DiI into the fourth mammary fat pad or spontaneous tumor of FVB control and MMTV-PyMT mice at 6, 9, and 12 weeks of age, and harvested DRG from each spinal level (**Fig. 1a**). We observed similarly low levels of DiI-labeled DRG neurons in both FVB and MMTV-PyMT mice at 6 weeks of age, suggestive of baseline mammary tissue innervation (**Fig. 1b**). At 9 and 12 weeks of age, during early and late stages of tumor development, MMTV-PyMT mice exhibited a significant increase in DiI-labeled neurons compared to FVB controls. There was no significant difference in innervation between 9- and 12-week MMTV-PyMT mice, indicating that sensory nerve recruitment occurs early during tumor progression^29^. We identified T8 to L2 DRGs as the predominant source of sensory innervation into abdominal mammary tumors, which corresponds anatomically to the spinal levels that typically supply sensory input to the lower mammary glands (**Fig. 1c**)^30^.

**Figure 1.**
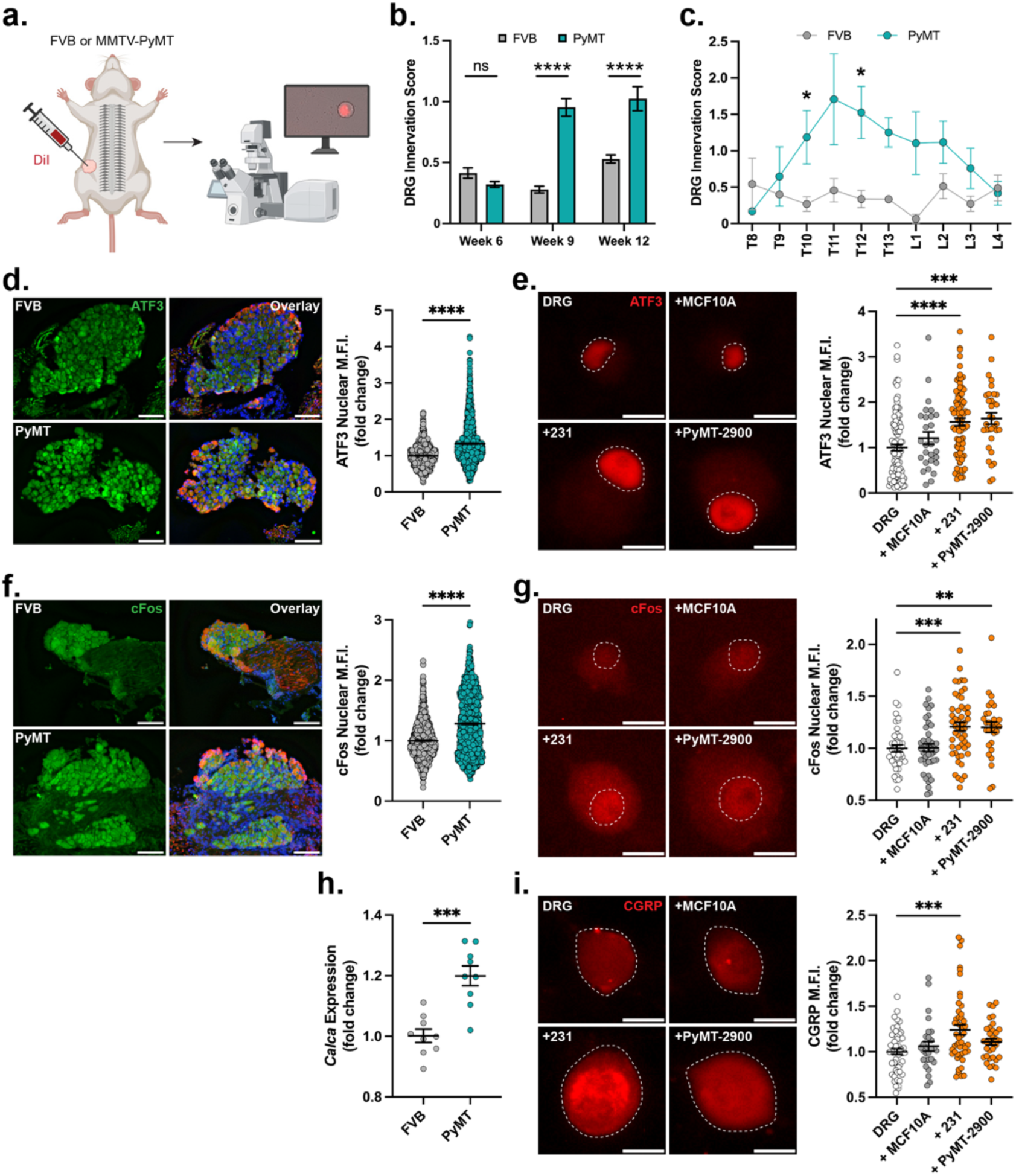
TNBC promotes increased innervation and activation in dorsal root ganglia neurons. **a)** Schematic of retrograde tracing experiment. Lipophilic dye DiI was injected into the fourth right mammary fat pad or tumor of FVB or MMTV-PyMT (PyMT) mice at 6, 9, and 12 weeks of age (n=3 mice/group; n=2 for MMTV-PyMT week 12). Three days later, DRG were harvested from all spinal levels and plated for analysis of DiI-labeled neurons by confocal microscopy. **b)** Quantification of DRG innervation score across all spinal levels, with each data point representing one spinal level per mouse (FVB: week 6, n=34; week 9, n=31; week 12, n=38; PyMT: week 6, n=45; week 9, n=36; week 12, n=29). **c)** DRG innervation score by spinal level at week 9. **d)** Nuclear ATF3 mean fluorescence intensity (MFI) in T12-L1 DRG sections from 13-week-old FVB (n=3 mice, 967 neurons) or MMTV-PyMT (n=3 mice, 1637 neurons) mice. DRG stained for DAPI (blue), β3-tubulin (red), and ATF3 (green). Scale, 100 µm. **e)** Nuclear ATF3 MFI in DRG neurons from C57BL/6J mice cultured alone (n=5 biological replicates, 107 neurons) or co-cultured with MCF10A cells (n=3, 28 neurons), human MDA-MB-231 (231) cells (n=5, 81 neurons), or murine PyMT-2900 cells (n=3, 33 neurons) for 12 days. Scale, 20 µm. **f)** Nuclear cFos MFI in DRG sections from 13-week-old FVB (n=3 mice, 1308 neurons) or MMTV-PyMT (n=3 mice, 1464 neurons) mice. DRG stained for DAPI (blue), β3-tubulin (red), and cFos (green). Scale, 100 µm. **g)** Nuclear cFos MFI in DRG neurons cultured alone (n=3, 44 neurons) or co-cultured with MCF10A cells (n=3, 43 neurons), 231 cells (n=3, 52 neurons), or PyMT-2900 cells (n=3, 30 neurons) for 8 days. Scale, 20 µm. **h)** *Calca* transcript levels in pooled T8-L4 DRG from 13-week-old FVB or MMTV-PyMT mice (n=3 mice/group, 3 technical replicates). **i)** CGRP MFI in DRG neuron cell bodies when cultured alone (n=3, 52 neurons) or co-cultured with MCF10A cells (n=3, 28 neurons), 231 cells (n=3, 50 neurons), or PyMT-2900 cells (n=3, 34 neurons) for 12 days. Scale, 20 µm. White dashed line indicates neuronal nuclei (e, g) or cell body (I). Data presented as mean ± SEM. Statistical significance determined by Student’s t-test (b, c, d, f, h) or one-way ANOVA (E, G, I), *p<0.05, **p<0.01, ***p<0.001, ****p<0.0001.

### Triple-negative breast cancer cells induce sensory neuron activation and stress response pathways

Neurite outgrowth is often preceded and accompanied by the activation of cellular stress or signaling pathways within neurons. To determine whether tumor cells trigger these neuronal responses, we assessed sensory neuron activation in both *in vivo* and *in vitro* models. DRG were harvested from mammary tumor-bearing MMTV-PyMT or control FVB mice for immunofluorescence staining or RNA isolation. In addition, DRG neurons from C57BL/6J mice were dissociated and cultured alone or co-cultured with non-tumorigenic MCF10A breast epithelial cells, MDA-MB-231 (231) human TNBC cells, or PyMT-2900 and PY8119 murine TNBC cells.

We performed immunofluorescence staining for nuclear ATF3 and cFos, two transcription factors upregulated in response to cellular stress and neuronal activity. ATF3 is a marker of nerve injury and is associated with regenerative programs, including neurite outgrowth, while cFos is an immediate early gene (IEG) rapidly induced following neuronal activation and calcium influx^31–35^. DRG from tumor-bearing MMTV-PyMT mice exhibited a significant increase in nuclear ATF3 staining compared to FVB controls (**Fig. 1d**). *In vitro*, DRG neurons co-cultured with human or murine TNBC cells, but not MCF10A cells, showed significantly elevated nuclear ATF3 expression compared to DRG cultured alone (**Fig. 1e, S1a, S2a**). Consistent with an activation response, nuclear cFos expression was increased in DRG from MMTV-PyMT mice relative to FVB controls, and co-culture with TNBC cell lines, but not MCF10A cells, led to a significant increase in neuronal cFos expression (**Fig. 1f, 1g, S1b, S2b**).

Previous studies have shown that calcitonin gene-related peptide (CGRP)-expressing nociceptive neurons innervate human and murine TNBC tumors^16^. CGRP, a neuropeptide involved in pain signaling and neurogenic inflammation, is known to increase following neuronal activation or injury^36–39^. We observed a significant increase in the expression of *Calca*, the gene encoding CGRP, in DRG from MMTV-PyMT mice relative to FVB controls (**Fig. 1H**). *In vitro*, CGRP expression was significantly increased in DRG neurons co-cultured with 231 cells, but not with murine TNBC cell lines (**Fig. 1I, S1C, S2C**). In summary, TNBC tumors broadly stimulate stress- and activation-associated pathways in sensory neurons *in vitro* and *in vivo*.

### Sensory neuron activation by tumor cells is partially mediated by TRPV1

TRPV1 is non-selective cation channel that, upon stimulation, triggers sensory neuron activation through calcium influx and promotes neurite outgrowth^23–25^. Although TRPV1-expressing nociceptive neurons are found in human TNBC tumors and high TRPV1 expression is associated with decreased patient survival, its role in tumor-nerve crosstalk has not been directly studied^15^. TRPV1 activation drives cFos signaling and increases CGRP expression^37, 39–41^. Given the observed effects of TNBC cells on these activation markers (**Fig. 1**), we hypothesized that tumor cells stimulate TRPV1 signaling, leading to neuronal activation and potentially promoting neurite outgrowth.

First, we measured TRPV1 expression in DRG neurons cultured alone or co-cultured with MCF10A or TNBC cells. While MCF10A cells induced a significant increase in TRPV1 expression, co-culture with 231 cells led to a more robust response (**Fig. 2a, S1d**). DRG co-cultured with PY8119 cells also exhibited a significant increase in TRPV1 expression, while PyMT-2900 co-culture did not elicit a significant change (**Fig. 2a, S1d, S2d**). Since TRPV1 expression levels do not necessarily reflect its activation status, we next sought to functionally test the involvement of TRPV1 signaling in sensory neuron activation^42^. In contrast to DRG from C57BL/6J mice, DRG isolated from *Trpv1*^-/-^ mice co-cultured with 231 cells did not exhibit a significant increase in cFos or CGRP expression (**Fig. 2b, 2c, S3**).

**Figure 2.**
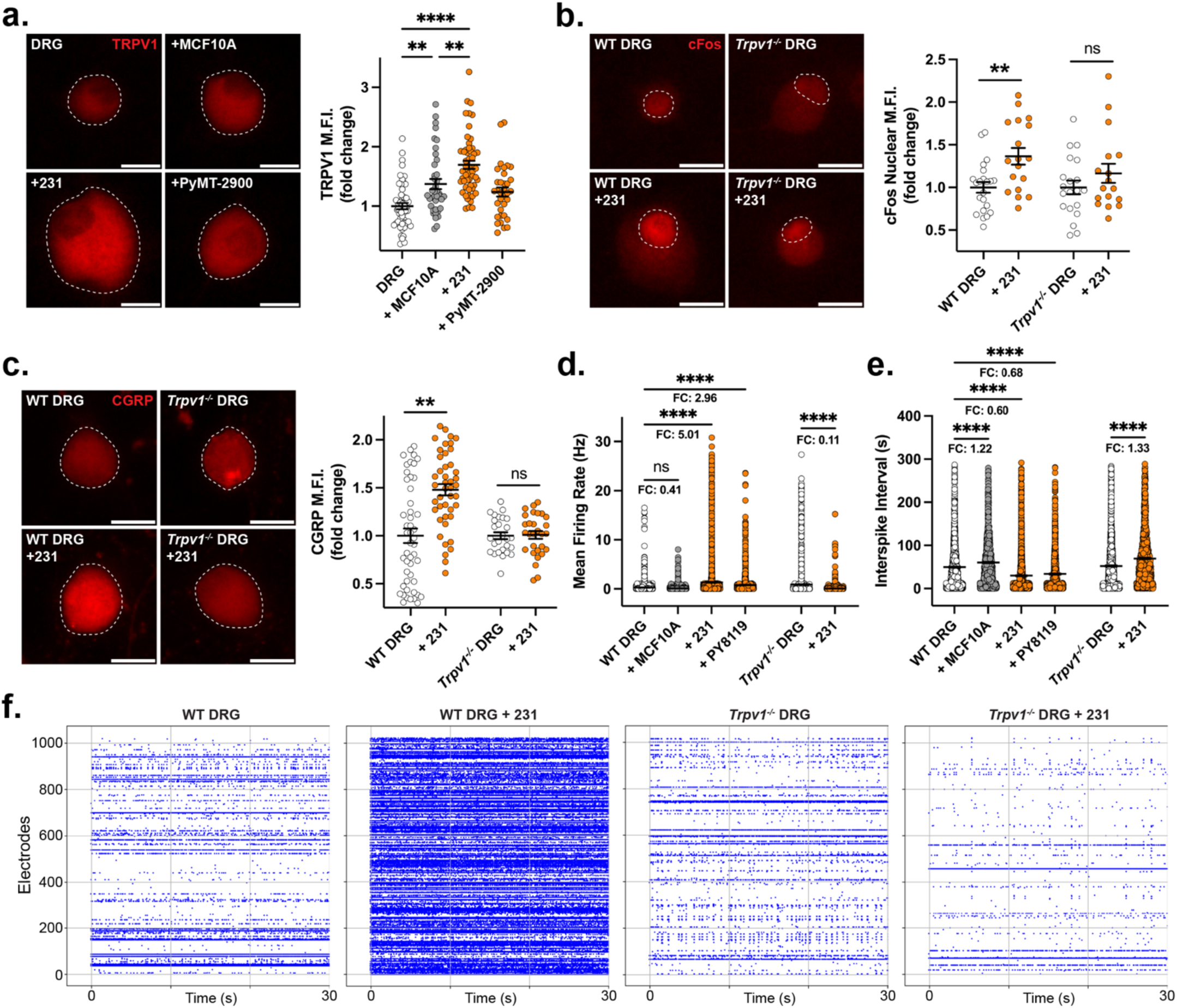
TNBC cells activate sensory neurons through TRPV1. **a)** TRPV1 mean fluorescence intensity (MFI) in DRG neuron cell bodies from C57BL/6J mice cultured alone (n=3 biological replicates, 52 neurons) or co-cultured with MCF10A cells (n=3, 39 neurons), MDA-MB-231 (231) cells (n=3, 54 neurons), or PyMT-2900 cells (n=3, 34 neurons) for 12 days. Scale, 20 µm. **b)** Nuclear cFos MFI in DRG neurons from C57BL/6J (WT) or *Trpv1^-/-^* mice cultured alone (WT: n=2, 22 neurons; *Trpv1^-/-^*: n=2, 20 neurons) or co-cultured with 231 cells (WT: n=2, 18 neurons; *Trpv1^-/-^*: n=2, 17 neurons) for 8 days. Scale, 20 µm. **c)** CGRP MFI in DRG neuron cell bodies from C57BL/6J (WT) or *Trpv1^-/-^* mice cultured alone (WT: n=49, 22 neurons; *Trpv1^-/-^*: n=2, 27 neurons) or co-cultured with 231 cells (WT: n=3, 44 neurons; *Trpv1^-/-^*: n=2, 29 neurons) for 12 days. Scale, 20 µm. White dashed line indicates neuronal cell body (a, c) or nuclei (b). **d-e)** DRG were cultured for 21 days on microelectrode arrays (MEAs) and bioelectric activity was measured for 300 seconds at endpoint. DRG from SF-W (WT) mice were cultured alone (n=4) or co-cultured with MCF10A (n=3), 231 (n=4), or PY8119 cells (n=2). DRG from *Trpv1^-/-^* mice were cultured alone (n=3) or co-cultured with 231 cells (n=3). **d)** Mean firing rate is the number of spikes per second per electrode (Hz). **e)** Interspike interval is plotted as the average time between two spikes per electrode. **f)** Representative raster plots, with each data point representing a detectable spike within the 30-second window. Data presented as mean ± SEM. Statistical significance determined by Student’s t-test (b, c, d, e) or one-way ANOVA (a, d, e), *p<0.05, **p<0.01, ***p<0.001, ****p<0.0001.

Since TRPV1 activation leads to calcium influx and can initiate action potentials in sensory neurons, we used microelectrode arrays (MEAs) to assess whether TNBC cells stimulate neuronal activity via TRPV1. DRG from female WT or *Trpv1^-/-^* mice were cultured alone or co-cultured with MCF10A or TNBC cells for 21 days on MEAs after which spontaneous neuronal activity was recorded over a 5-minute period. Co-culture with 231 and PY8119 cells, but not MCF10A cells, significantly elevated the mean firing rate in WT DRG, while MCF10A cell co-culture decreased firing (**Fig. 2d, 2e, S4a**). In contrast, *Trpv1^-/-^* DRG co-cultured with 231 cells showed a significant decrease in mean firing rate. Next, we assessed the interspike interval, where a shorter interval reflects increased frequency of neuronal firing. WT DRG co-cultured with 231 and PY8119 cells significantly decreased the interspike interval, whereas MCF10A co-culture led to an increase (**Fig. 2e, 2f, S4a**). *Trpv1^-/-^* DRG co-cultured with 231 cells exhibited an increase in time between spikes. WT DRG cultured with conditioned media (CM) from 231 cells led to varied effects, with no significant increase in mean firing rate but a significant decrease in interspike interval (**Fig. S4a, S4c-d**). Together, these data show evidence that TNBC cells drive increased electrical activity in DRG neurons in a TRPV1-dependent manner, raising the possibility that TRPV1 also promotes neurite outgrowth.

### Triple-negative breast cancer cells drive neurite outgrowth in a TRPV1-dependent manner

We next sought to determine whether TNBC cells drive sensory neuron growth *in vitro* and if these effects are dependent on neuronal TRPV1. We measured the effect of tumor cells or their conditioned media (CM) on morphological features linked to neurite initiation, growth, and maturation. WT DRG cultured with non-tumorigenic MCF10A cell CM or MCF10A cells induced no significant changes in neuron morphology (**Fig. 3c-f, S5a-e**). In contrast, 231 cell CM or 231 cell co-culture significantly enhanced several hallmarks of neuronal growth (**Fig. 3, S5a-e**). WT DRG exhibited a significant increase in the number of primary neurites extending from neuron somas when cultured with 231 cell CM or 231 cells, indicative of enhanced neuritogenesis (**Fig. 3c, S5b**)^43^. Additionally, 231 cell CM and 231 cell co-culture significantly increased both the maximum diameter of neurites and the area of neuron cell bodies, features associated with neuronal growth and regeneration (**Fig. 3d-e, S5c-d**)^44–46^. Direct co-culture with 231 cells, but not 231 cell CM, significantly increased neurite bundling, a process known as fasciculation that facilitates coordinated axon growth during regeneration and development (**Fig. 3a, 3f, S5e**)^47^. In DRG from *Trpv1^-/-^* mice, however, 231 cell CM and 231 cell co-culture failed to induce significant changes in primary neurite number and neurite diameter (**Fig. 3b-d, S5a-c**). A modest increase in neuron cell body area and neurite fasciculation was observed in *Trpv1^-/-^* neurons co-cultured with 231 cell CM or cells, but these effects were attenuated compared to WT neurons (**Fig. 3e, S5d-e**).

**Figure 3.**
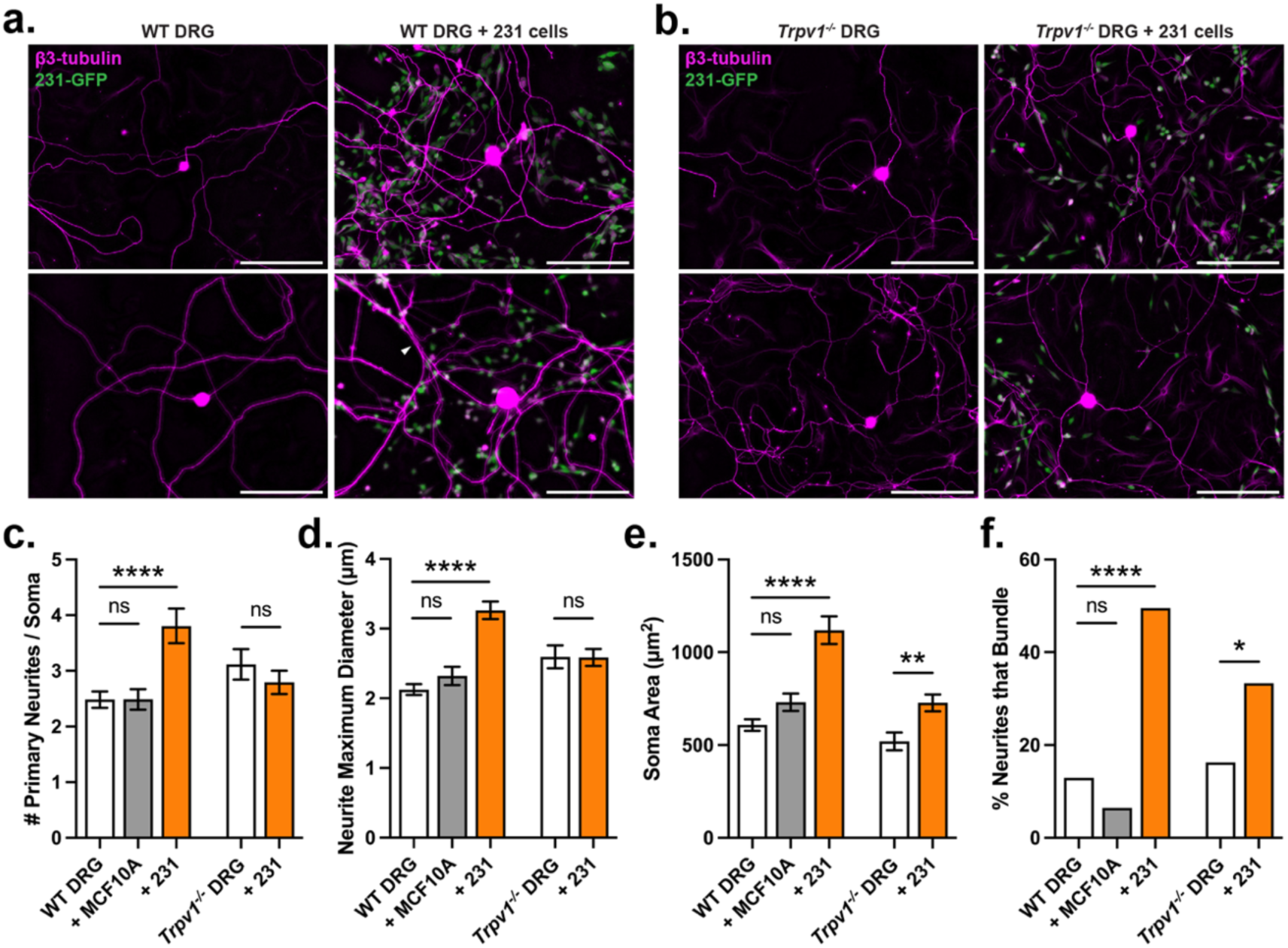
TNBC cells promote sensory neurite outgrowth in 2D in a TRPV1-dependent manner. **a-b)** Representative images of (**a**) C57BL/6J (WT) or (**b**) *Trpv1^-/-^*DRG cultured alone or co-cultured with GFP+ MDA-MB-231 (231) cells for 12 days and stained for β3-tubulin. White arrow indicates neurite fasciculation. Scale bar, 200 μm. **c-f)** DRG were analyzed for various morphological features linked to neurite initiation and growth by ImageJ. **c)** Number of primary neurites per soma. WT DRG cultured alone (n=8 biological replicates, 56 neurons) or co-cultured with MCF10A cells (n=4, 31 neurons) or 231 cells (n=4, 31 neurons). *Trpv1^-/-^* DRG cultured alone (n=4, 26 neurons) or co-cultured with 231 cells (n=4, 29 neurons). **d)** Neurite maximum diameter per field of view (FOV). WT DRG cultured alone (n=6, 39 FOV) or co-cultured with MCF10A cells (n=2, 16 FOV) or 231 cells (n=4, 28 FOV). *Trpv1^-/-^*DRG cultured alone (n=4, 26 FOV) or co-cultured with 231 cells (n=4, 28 FOV). **e)** Soma area. WT DRG cultured alone (n=8, 59 neurons) or co-cultured with MCF10A cells (n=4, 31 neurons) or 231 cells (n=4, 29 neurons). *Trpv1^-/-^* DRG cultured alone (n=4, 26 neurons) or co-cultured with 231 cells (n=4, 29 neurons). **f)** Percent of neurites that bundle. WT DRG cultured alone (n=8, 18/139) or co-cultured with MCF10A cells (n=4, 5/77) or 231 cells (n=4, 27/81). *Trpv1^-/-^*DRG cultured alone (n=4, 13/80) or co-cultured with 231 cells (n=4, 27/81). Data presented as mean ± SEM. Statistical significance determined by one-way ANOVA (c, d, e) or Fisher’s exact test (f), *p<0.05, **p<0.01, ***p<0.001, ****p<0.0001.

While 2D assays provide valuable insights into cancer-nerve interactions, they lack the ECM components that accurately mimic the physiological environment. Complex mammary tissues can grow in 3D hydrogel scaffolds composed of laminin, hyaluronan, fibronectin, and collagen^48, 49^. We confirmed that these 3D hydrogels support the growth of DRG explants, which extend long neurites over a 16-day period (**Fig. S6**). Next, to investigate whether TNBC cell secretions drive directional outgrowth of neurites, DRG from WT or *Trpv1^-/-^* mice were positioned in the center of a hydrogel, with 231 cells placed on one side and no cells placed on the other side (**Fig. 4a**). DRG from WT mice exhibited a significant increase in neurite density towards 231 cells that was abrogated with *Trpv1^-/-^* DRG (**Fig. 4b-c**). Together, these results demonstrate that tumor cells induce significant morphological changes in sensory neurons associated with increased neuronal process outgrowth and innervation.

**Figure 4.**
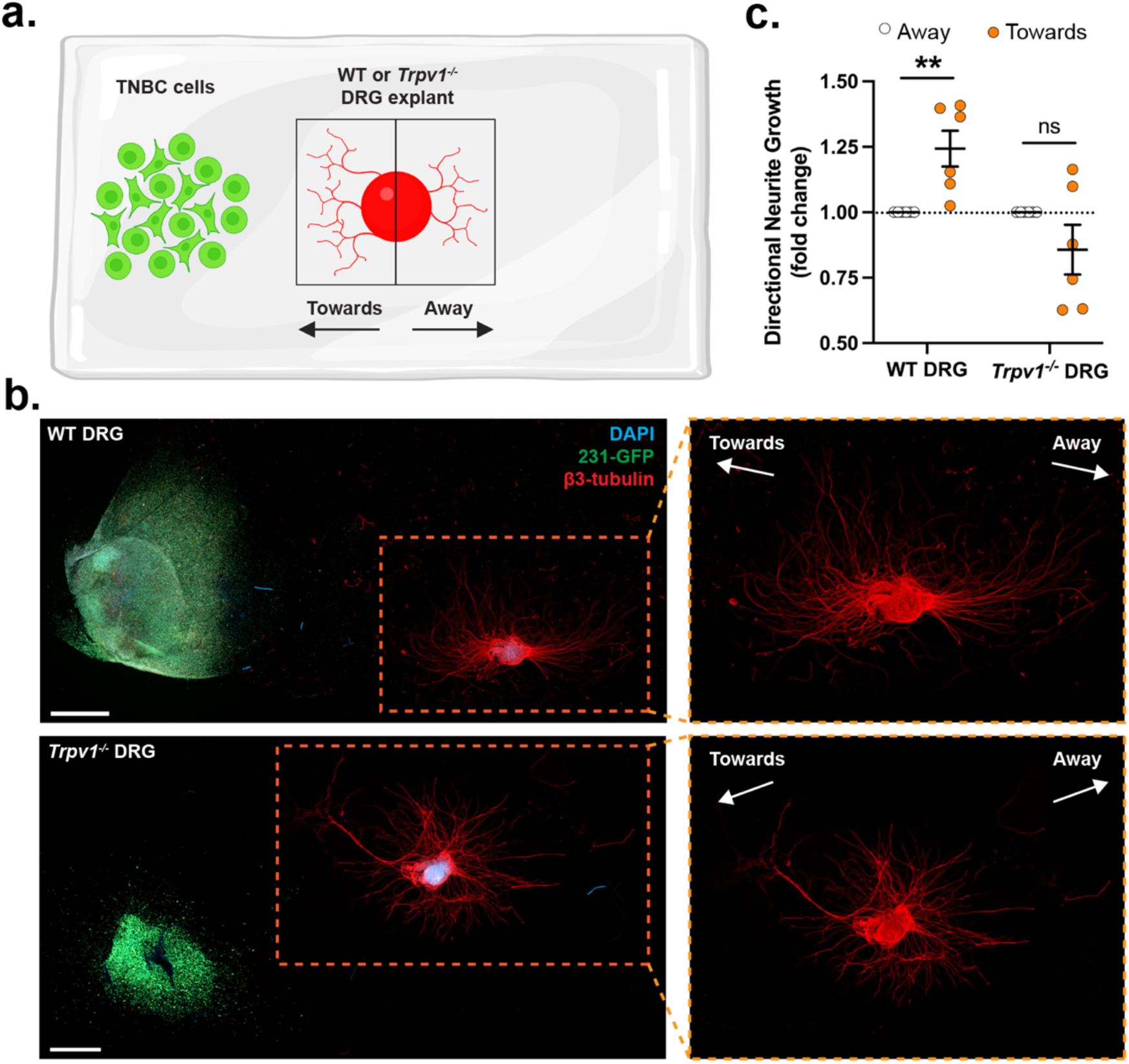
TNBC cells promote directional neurite outgrowth from DRG explants in a TRPV1-dependent manner. **a)** Schematic of 3D neurite directionality experiment. DRG explants from C57BL/6J (n=6 gels) or *Trpv1^-/-^* (n=6 gels) mice were embedded in the center of a 3D hydrogel, with GFP+ MDA-MB-231 (231) cells on the left side. Gels were fixed after 16-20 days in culture and stained for β3 tubulin and DAPI. ImageJ was used to quantify β3-tubulin-positive area in the direction towards or away from 231 cells. **b)** Representative images of DRG explants after 16 days in culture. Scale bar, 1000 μm. **c)** Quantification of β3-tubulin signal towards 231 cells, graphed as fold change relative to the signal away from 231 cells within the same gel. Data presented as mean ± SEM. Statistical significance determined by student’s T-test, **p<0.01.

### IL-6 acts downstream of TRPV1 to promote neurite outgrowth in response to triple-negative breast cancer cells

To elucidate the mechanism by which TRPV1 activation promotes neurite outgrowth, we analyzed RNA sequencing data from DRG cultured alone or co-cultured with 231 cells. The sequencing was performed and made publicly available when we investigated the effect of DRG on TNBC cells, but was not analyzed^15^.

Gene ontology pathways analysis revealed that co-culture with 231 cells led to an upregulation of genes involved in cell proliferation, cell migration, and developmental processes in DRG, suggesting activation of pathways associated with neurite outgrowth (**Fig. 5a**). TRPV1 stimulation induces calcium influx, which leads to the activation of various transcription factors and downstream effects^40, 50^. To identify candidate pathways involved in neurite outgrowth, we performed transcription factor enrichment analysis on genes upregulated in DRG co-cultured with 231 cells. We found that cJun was among the most highly enriched transcription factors (**Fig. 5b**). cJun is activated downstream of TRPV1 stimulation and promotes neurite outgrowth by regulating a broad set of genes, making it a strong candidate for further analysis^40, 51–54^. Specifically, Interleukin 6 (*Il6)*, a known cJun target gene and key mediator of neurite outgrowth, was highly upregulated, showing a 5.27-fold increase in co-culture compared to DRG alone (**Fig. 5c**)^55–59^. Based on these data, we hypothesized that TNBC cells activate TRPV1 on sensory neurons, triggering cJun activation and subsequent *Il6* expression, which promotes neurite outgrowth (**Fig. 5d**).

**Figure 5.**
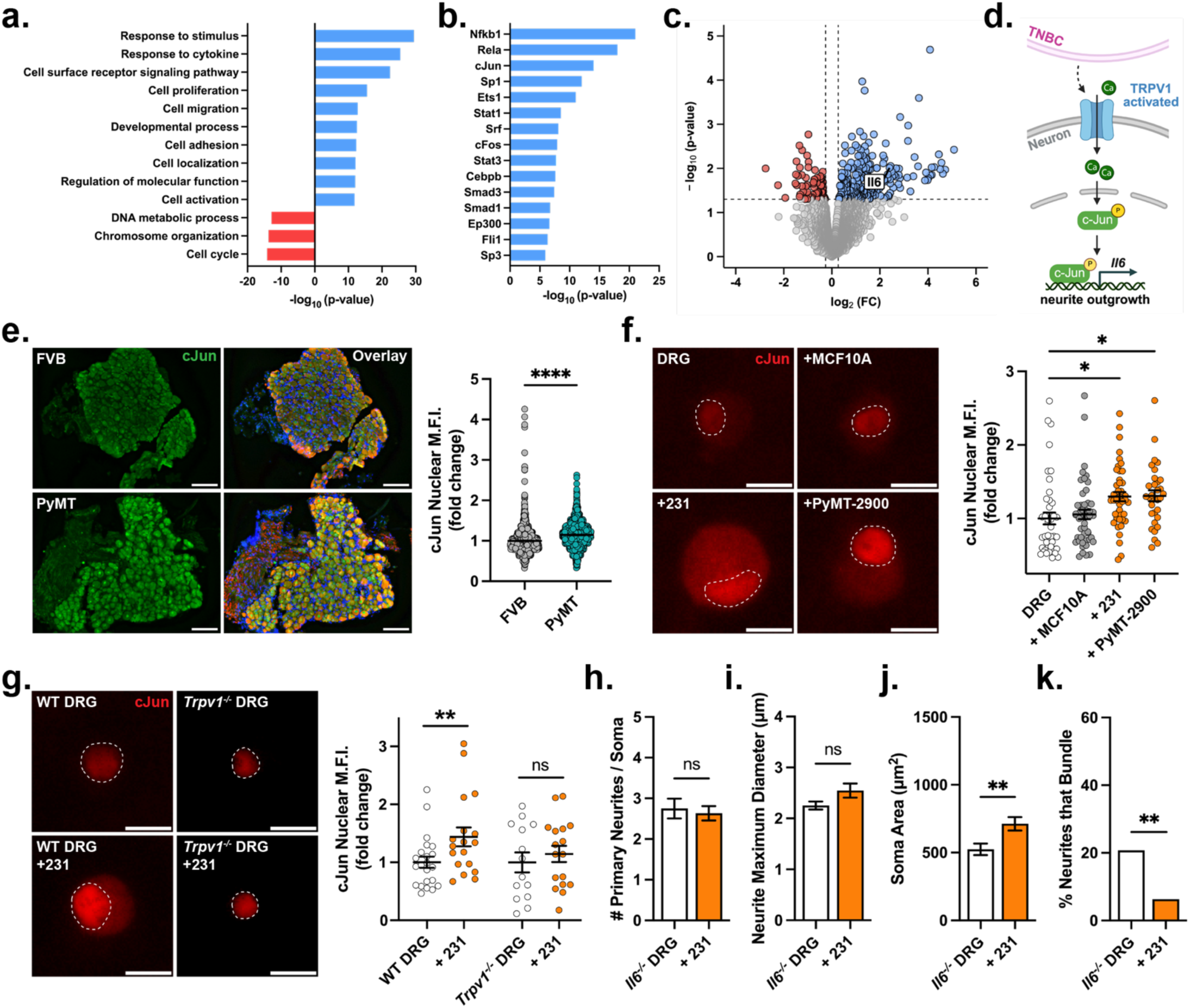
IL-6 acts downstream of TRPV1 to promote neurite outgrowth in response to TNBC cells. **a)** Gene ontology (GO) biological processes enriched among genes upregulated in WT DRG co-cultured with MDA-MB-231 (231) TNBC cells compared to WT DRG cultured alone. Blue bars indicate positively enriched pathways, red bars indicate negatively enriched pathways. **b)** Transcription factors enrichment analysis of genes upregulated in WT DRG co-cultured with 231 cells. **c)** Volcano plot of differentially expressed genes in WT DRG co-cultured with 231 cells vs. WT DRG alone. Genes significantly upregulated are shown in bule, downregulated in red. **d)** Nuclear cJun mean fluorescence intensity (MFI) in T12-L1 DRG sections from 13-week-old FVB (n=3 mice, 1570 neurons) or MMTV-PyMT (n=3 mice, 1924 neurons) mice. DRG stained for DAPI (blue), β3-tubulin (red), and cJun (green). Scale, 100 µm. **e)** Nuclear cJun MFI in DRG neurons cultured alone (n=3 biological replicates, 43 neurons) or co-cultured with MCF10A cells (n=3, 48 neurons), 231 cells (n=3, 44 neurons), or PyMT-2900 cells (n=3, 32 neurons) for 4 days. Scale, 20 µm. **f)** Nuclear cJun MFI in DRG neurons from C57BL/6J (WT) or *Trpv1^-/-^* mice cultured alone (WT: n=2, 22 neurons; *Trpv1^-/-^*: n=2, 20 neurons) or co-cultured with 231 cells (WT: n=2, 18 neurons; *Trpv1^-/-^*: n=2, 17 neurons) for 4 days. Scale, 20 µm. **g-j)** *Il6^-/-^* DRG cultured alone or co-cultured with 231 cells and stained for β3-tubulin. **g)** Number of primary neurites per soma. *Il6^-/-^* DRG cultured alone (n=3 biological replicates, 28 neurons) or co-cultured with 231 cells (n=4, 30 neurons). **h)** Neurite maximum diameter per field of view (FOV). *Il6^-/-^* DRG cultured alone (n=2, 16 neurons) or co-cultured with 231 cells (n=2, 16 neurons). **i)** Soma area. *Il6^-/-^* DRG cultured alone (n=3, 28 neurons) or co-cultured with 231 cells (n=4, 30 neurons). **j)** Percent of neurites that bundle. *Il6^-/-^* DRG cultured alone (n=3, 16/77) or co-cultured with 231 cells (n=4, 5/79). Data presented as mean ± SEM. Statistical significance determined by Student’s t-test (d, f), one-way ANOVA (e, g, h, i), or Fisher’s exact test (j), *p<0.05, **p<0.01, ***p<0.001, ****p<0.0001.

We validated the increased cJun activity observed in our RNAseq data using immunofluorescence *in vivo* and *in vitro*. Nuclear cJun expression was significantly elevated in DRG sections from tumor-bearing MMTV-PyMT mice compared to FVB controls (**Fig. 5e**). Moreover, co-culture of DRG neurons with TNBC cells, but not MCF10A cells, led to a significant increase in nuclear cJun expression (**Fig. 5f, S7a-b**). Co-culture of 231 cells with DRG neurons from *Trpv1^-/-^*mice did not result in a significant increase in cJun expression, suggesting that TRPV1 signaling is required for tumor cell-driven upregulation of cJun (**Fig. 5g, S7c**).

To investigate the role of IL-6 in promoting neurite outgrowth, DRG neurons from *Il6^-/-^* mice were cultured alone, with 231 cell CM, or with 231 cells, and morphological changes were assessed. Similar to the findings with *Trpv1^-/-^*DRG (**Fig. 3**), co-culture with 231 cell CM or 231 cells did not significantly alter neuritogenesis or neurite maximum diameter, while a slight increase in neuron cell body area was observed (**Fig. 5h-j, S8a-d**). In addition, 231 CM or 231 cells failed to induce an increase in neurite bundling in *Il6^-/-^* DRG neurons (**Fig. 5k, S8e**). Together, these findings suggest that TRPV1 activation promotes neurite outgrowth in part through cJun-mediated upregulation of IL-6 signaling.

### Knockout of Trpv1 slows tumor progression

To investigate the role of TRPV1 *in vivo*, murine PY8119 TNBC cells were injected into the fourth right mammary fat pads of 7-week-old C57BL/6J (WT) or *Trpv1^-/-^* mice. Loss of TRPV1 delayed tumor growth, with *Trpv1^-/-^* mice exhibiting significantly smaller tumor volumes at days 10 and 16 compared to WT mice (**Fig. 6a-b**). Tumor weight at the day 25 endpoint was significantly decreased in *Trpv1^-/-^* mice (**Fig. 6c**). Analysis of lung tissue with hematoxylin and eosin (H&E) staining found statistically significant decreases in both the number of lung metastasis and the percentage of mice with metastasis (**Fig. 6d-f**).

**Figure 6.**
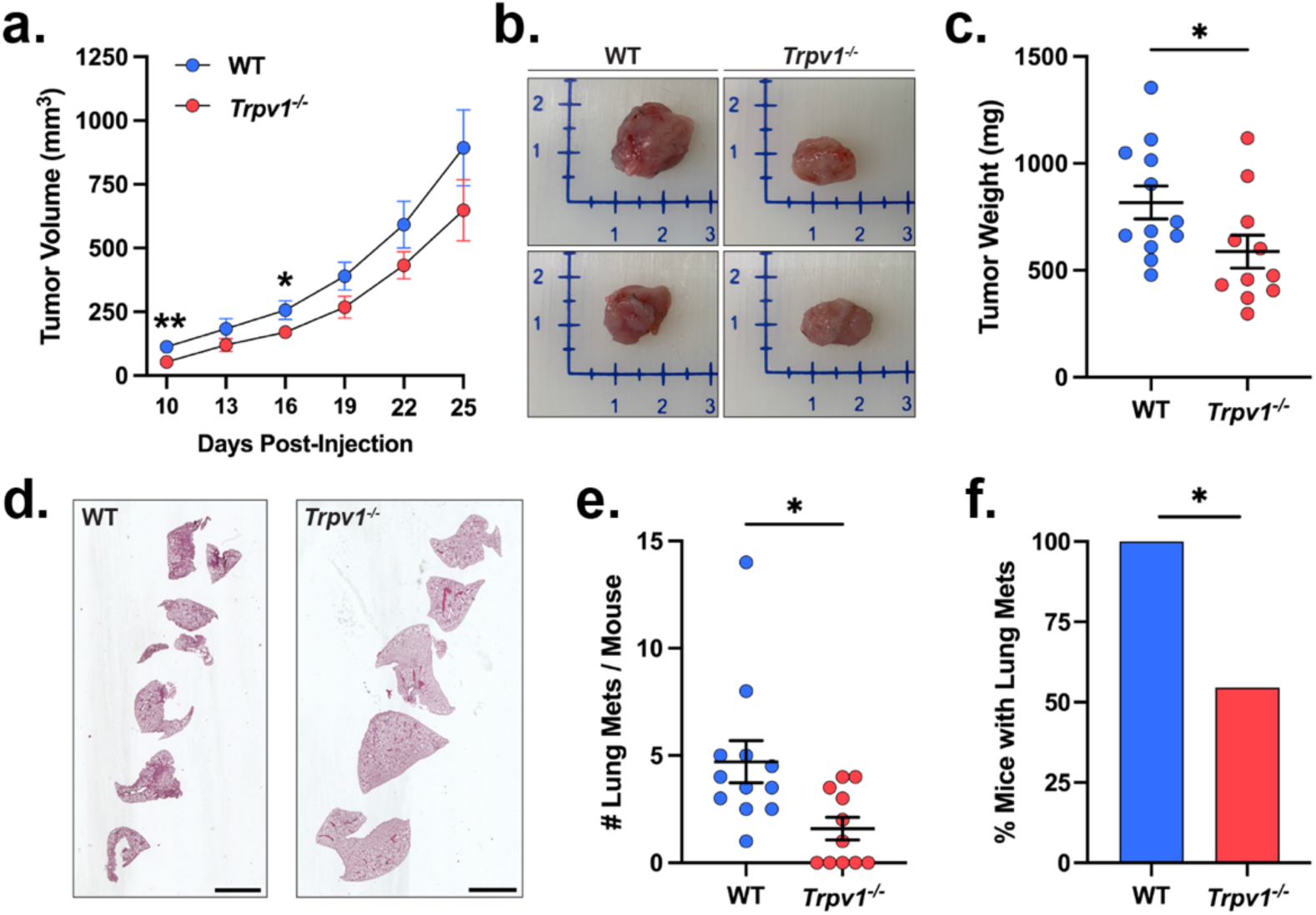
*Trpv1* knockout inhibits TNBC lung metastasis. Murine PY8119 TNBC cells were injected into the 4^th^ right mammary fat pads of C57BL/6J (WT, n=12) or *Trpv1^-/-^* mice (n=11). **a)** Tumor volume was measured using calipers every 3 days from when tumors were palpable. **b)** Representative images of tumors at day 25. **c)** Tumor weight at day 25. **d)** Representative sections of lung tissue stained with hematoxylin and eosin (H&E). Scale bar, 1000 μm. **e)** Number of lung metastases per mouse. **f)** Percent mice with lung metastasis. Data presented as mean ± SEM. Statistical significance determined by student’s T-test (a, c, e) or Fisher’s exact test (f), *p<0.05.

## Discussion

Despite growing evidence linking nerves to tumor progression, the mechanisms that drive tumor innervation remain poorly understood. Given that TNBC is predominantly innervated by sensory nerves, we asked whether tumor-derived cues might activate sensory neurons and thereby promote neurite outgrowth into the tumor microenvironment. We find that TNBC cells activate sensory neurons via TRPV1, leading to increased expression of activity markers, elevated bioelectric activity, and enhanced neurite outgrowth in both 2D and 3D models. These effects are mediated through TRPV1-dependent c-Jun signaling and downstream IL-6. Lastly, we show that knockout of TRPV1 in sensory nerves decreases tumor growth and metastasis to the lung. This is the first comprehensive description using both 2D and 3D models of tumor cell-driven effects on sensory nerve activation, electrical activity, and growth. Our results identify novel strategies that could suppress innervation and thereby reduce the pro-tumorigenic effects of nerves.

First, we find that TNBC cells activate sensory neurons via TRPV1, resulting in increased expression of activity markers and enhanced bioelectric activity, features known to be associated with nerve regeneration in the context of injury. Tumor-bearing MMTV-PyMT mice demonstrated an increase in cFos and CGRP in their DRG compared to FVB controls. These effects were validated in *in vitro* cultures and were abrogated in *Trpv1* knockout neurons. Using MEAs, we showed that co-culture with TNBC cells increases the firing rate and decreases the interspike interval of DRG neurons. To our knowledge, this is the first evidence of breast tumor cells driving bioelectric activity in sensory neurons. Increased sensory nerve activation and firing has been well described in the context of nerve injury and regeneration, where DRGs upregulate transcriptional programs that drive growth and fire in a bursting pattern^60^. These data are perhaps not surprising given that the tumor microenvironment has long been described as a wound that never heals, creating an environment akin to nerve injury which can drive regeneration. Tumor growth is also associated with increased blood vessel density, which often co-occurs with sensory nerve growth^61^. Further, while synaptic structures have not been described between sensory neurons and breast cancer cells in the primary tumor, recent studies have identified functional neuron-to-tumor synapses in brain metastases and gliomas, where neuronal activity promotes tumor growth through glutamatergic signaling^62–64^. Small cell lung cancer cells co-cultured with neurons exhibit synaptic gene expression and electrophysiological responses^65^. These findings raise the possibility that certain cancers, including peripheral tumors, may engage neurons through more structured, contact-dependent mechanisms in addition to paracrine signaling. Future studies are needed to dissect whether TNBC-driven effects on sensory nerve activation and electrical activity are mediated by actual synapses or combinations of local pro-regenerative cues that activate sensory neurons and promote outgrowth.

Our data demonstrate that some effects of tumor cells on nerves are driven by direct tumor cell-nerve contact, but some are not. Indeed, in 2D cell culture assays, both conditioned media from TNBC cells and direct co-culture enhanced several morphological features associated with neurite outgrowth, including increased neuritogenesis, greater neurite diameter, and enlarged soma size. Interestingly, direct co-culture, but not conditioned media, increased neurite fasciculation. This bundling was also observed in *Trpv1* knockout DRG co-cultured with 231 cells, suggesting that tumor cell-driven fasciculation is a contact-dependent, TRPV1-independent process. It may instead be mediated by direct tumor cell-neuron contact via adhesion molecules such as integrins or cadherins, which could in turn upregulate adhesion molecule and receptor expression in neurons^66^. Alternatively, tumor-induced stiffening of the extracellular matrix or spatial constraints imposed by tumor cells may restrict neurite pathfinding and promote bundling^47^. In 3D hydrogels, where TNBC cells and DRG explants are physically separated, directional neurite outgrowth towards tumor cells was observed in wild-type neurons alone, supporting a model in which secreted factors drive TRPV1-dependent neurite extension. Additionally, this 3D system offers a more physiologically relevant platform to study tumor–neuron interactions, preserving the architecture of the ganglia and supporting endogenous cell-cell signaling that may be lost in dissociated cultures. Importantly, neuronal cell bodies remain spatially distinct from the tumor cells, consistent with the *in vivo* organization in which peripheral ganglia innervate tumors without migrating into them. Interestingly, conditioned media had varied effects on the bioelectric activity of DRG neurons, with no change in mean firing rate but a decrease in interspike interval. This suggests that CM may lead to alterations in DRG firing patterns that differ from direct co-culture. The tumor microenvironment provides many chemical and biophysical cues that are known to activate TRPV1 on sensory neurons, such as heat (>43°C), mild acidification, inflammatory mediators, and NGF^5, 26, 27^.One possibility is that direct tumor-neuron contact allows for additional modes of interaction beyond TRPV1 activation, such as mechanically gated channels, ephaptic coupling, or even synapse-like formation^67, 68^. Further, it is possible that sustained tumor-cell nerve interactions over time are needed to produce the soluble cues needed for these effects to be seen. Most studies to date have focused on tumor-secreted neurotrophic factors, such as NGF and BDNF, or axon guidance molecules, such as SLIT2, semaphorins, or ephrin-B1^3, 8, 9, 16, 21, 69–75^. While these signals are critical for neurogenesis during development and regeneration, other regulatory mechanisms, including physical properties of the ECM, cell-cell adhesions, inflammatory cytokines and injury signaling, and notably, neuronal activation, also contribute to neurite outgrowth^76–79^. While it is likely that a combination of TME cues contribute to the sensory neuron activation we describe here, future studies are needed to dissect the contribution of individual signals to sensory innervation.

Lastly, we observed a critical role of TRPV1 in tumor cell-driven neuron activation and neurite outgrowth, and observed reduced tumor growth and metastasis in *Trpv1* knockout mice. These results suggest that inhibiting TRPV1 could be a viable strategy to disrupt tumor–nerve interactions.TRPV1 antagonists were originally developed for the treatment of chronic pain and inflammatory conditions due to the channel’s established role in nociception^80, 81^. Several potent TRPV1 inhibitors, including AMG 517, SB-705498, and JNJ-17203212, showed promising analgesic activity in preclinical models and entered early-phase clinical trials^82, 83^. However, development was largely halted due to adverse effects, including hyperthermia and impaired noxious heat sensation, which raised safety concerns about systemic TRPV1 blockade^84, 85^. More recently, new approaches have been developed to overcome the limitations of earlier TRPV1 inhibitors, including modality-selective antagonists that block specific modes of TRPV1 activation while sparing thermosensation, peripherally restricted compounds that minimize central side effects, and localized delivery strategies that reduce systemic exposure. Continued development of safer and more targeted TRPV1 inhibitors, alongside efforts to identify the specific tumor-derived cues that activate TRPV1, may pave the way for novel therapeutic strategies that disrupt nerve-cancer interactions in TNBC and beyond.

## Methods

### Animal care

All animal procedures were reviewed and approved by the Tufts University Institutional Animal Care and Use Committee. All mice were housed 3-5 mice per cage in an on-site housing facility with access to standard food, water, and 12/12 light cycle.

### Dorsal root ganglia dissection

Dorsal root ganglia (DRG) dissections were performed as previously described^15, 86^. Briefly, mice were euthanized by CO_2_ and spinal columns were isolated. Lateral cuts were made along the spinal column to expose the spinal cord. Ganglia that lie adjacent to the spinal cord were collected using sterile tweezers. DRG were either fixed in 4% paraformaldehyde and embedded in paraffin for immunofluorescence imaging, collected for RNA extraction, dissociated into single cell suspension for 2D plating, or cultured as explants in 3D hydrogels, as explained below.

### Retrograde tracing

Lipophilic tracer 1,1’-Dioctadecyl-3,3,3’,3’-Tetramethylindocarbocyanine Perchlorate (DiI, Cat. No. D3911, Invitrogen) was prepared using 100% ethanol for a concentration of 2.5 mg/ml. At 6, 9, and 12 weeks of age, 25 μl DiI was injected into the 4^th^ right mammary fat pad or corresponding tumor of three female FVB (Strain #001800, Jackson Laboratory) or PyMT-MMTV (Strain #002374, Jackson Laboratory) mice. One of the week 12 MMTV-PyMT mice was euthanized prior to the planned endpoint in accordance with IACUC protocols due to tumor volume exceeding 2 cm^3^, and no data was collected from this animal. Three days after injection at each timepoint, mice were euthanized by CO_2_ and dorsal root ganglia were dissected individually from each spinal level, with each DRG placed in a separate tube for independent processing. Single cell dissociation of each DRG was performed with 1.25 mg/ml Collagenase A (Cat. No. 10103586001, Sigma) in HBSS for 1 hour at 37°C, followed by 1.2 mg/ml Trypsin (Cat. No. 25200114, Gibco) in HBSS for 15 min at 37°C. DRG were then mixed by gentle pipetting. Next, DRG were centrifuged at 1500 rpm for 3 min and resuspended in complete neurobasal media, which consists of: Neurobasal Medium (Cat. No. 21103-049, Gibco), 2% B27 Plus Supplement (Cat. No. A35828-01, Gibco), 1% GlutaMAX Supplement (Cat. No. 35050-061, Gibco), 1% Antibiotic-Antimycotic (Cat. No. 15240-062, Gibco), 25 ng/ml Recombinant Human β-NGF (Cat. No. 450-01, PeproTech), and 10 ng/ml Human GDNF Recombinant Protein (Cat. No. PHC7045, Gibco). 24-well glass bottom plates (Cat. No. P24-1.5H-N, Cellvis) were coated with 0.1 mg/ml Poly-D-Lysine (Cat. No. A38904-01) for 1 hour at 37°C and 20 μg/ml Laminin (Cat. No. L2020, Sigma) for 1 hour at 37°C. Cells from each dissociated DRG were seeded into its own well of a 24-well plate, maintaining one spinal level per well. In some cases, specific DRG could not be located during dissection and were therefore not included in subsequent analyses. 24-40 hours after plating, fluorescence and brightfield images were taken using the Zeiss Axio Observer (Zeiss) with six fields of view per well to visualize DiI-labeled neurons. Images were graded independently by two observers on a 0-4 scale based on the number and intensity of DiI-positive neurons. Scores from all fields and both reviewers were averaged to yield a single innervation score per DRG.

### Dorsal root ganglia immunofluorescence staining and qPCR

Three 13-week-old female FVB (Strain #001800, Jackson Laboratory) or PyMT-MMTV (Strain #002374, Jackson Laboratory) mice were euthanized and DRG were dissected from T10-L2 spinal levels. For immunofluorescence, DRG were fixed for 24 hours in 4% Paraformaldehyde (Cat. No. 15710-S, Electron Microscopy Sciences) in PBS, embedded in paraffin, and section into 10 μm sections. Sections were deparaffinized and antigen retrieval was performed in Citra Plus Solution (Cat. No. HK080, Biogenex). Sections were then blocked in PBS with 0.5% Tween-20 (Cat. No. P1379, Sigma), 10% Normal Donkey Serum (Cat. No. 017-000-121, Jackson ImmunoResearch), and 20 μg/mL AffiniPure Fab Fragment Donkey Anti-Mouse IgG (Cat. No. 715-007-003, Jackson ImmunoResearch), and incubated with primary antibodies (Table 1) overnight at 4°C. The following day, sections were incubated with 4’,6-diamidino-2-phenylindole (DAPI, dilution: 1/1000, Cat. No. D1306, Invitrogen) and fluorophore-conjugated donkey secondary antibodies (dilution: 1/500, Jackson ImmunoResearch), including Alexa Flour 488 and Alexa Flour 647, for 2 hours at room temperature. Imaging was performed using a Keyence BZ-X710 microscope at 20X magnification. Nuclear fluorescence intensity (ATF3, cFos, cJun) in DRG neurons was quantified using a custom ImageJ macro, with β3-tubulin identifying neurons and DAPI defining nuclear masks. Nuclear intensity values were normalized to β3-tubulin intensity to account for variances in staining efficiency.

**Table 1.** Primary antibodies and their dilutions used in this study.

| Target | Host | Company | Catalog # | DRG Section | 2D Culture | 3D Hydrogel |
| --- | --- | --- | --- | --- | --- | --- |
| β3-tubulin | Rabbit | Abcam | ab18207 | 1/200 | 1/500 | 1/500 |
| β3-tubulin | Mouse | Sigma-Aldrich | T8578 | | 1/500 | |
| cJun | Mouse | Santa Cruz | sc-74543 | 1/50 | 1/50 |  |
| cFos | Mouse | Santa Cruz | sc-8047 | 1/50 | 1/50 |  |
| ATF3 | Mouse | Santa Cruz | sc-518032 | 1/100 | 1/100 |  |
| CGRP | Goat | Abcam | ab36001 |  | 1/100 |  |
| TRPV1 | Rabbit | Alomone Labs | ACC-030 |  | 1/200 |  |
| E-cadherin | Mouse | Invitrogen | 13-1700 |  | 1/100 |  |

For quantitative PCR (qPCR), DRG from spinal levels T8-L4 were pooled from each mouse, and total RNA was extracted using a Quick-RNA Miniprep Kit (Cat. No. R1055, Zymo Research) according to the manufacturer’s instructions. Complementary DNA (cDNA) was synthesized using the High-Capacity cDNA Reverse Transcription Kit (Cat. No. 4368814, ThermoFisher Scientific) at the following thermocycler conditions: Step 1, 25°C for 10 min; Step 2, 37°C for 120 min; Step 3, 85°C for 5 min; Step 4, 4°C hold. qPCR was then performed using TaqMan Fast Advanced Master Mix (Cat. No. 4444557, ThermoFisher Scientific) and TaqMan Gene Expression Assays (Cat. No. 4453320, ThermoFisher Scientific) for mouse *Calca* (Assay ID: mm00801463_g1), with mouse *Gapdh* as the endogenous control (Assay ID: mm99999915_g1). Relative gene expression was calculated using the ΔΔCT method.

### Cell culture

MCF10A, MDA-MB-231, and PY8119 cells were obtained from ATCC. PyMT-2900 cells were a gift from Professor Richard Hynes’ lab at MIT. MCF10A cells were cultured in MEGM Mammary Epithelial Cell Growth Medium BulletKit (Cat. No. CC-3150, Lonza) without GA-1000 (gentamicin-amphotericin B) and supplemented with 100 ng/ml Cholera Toxin (Cat. No. C8052, Sigma). MDA-MB-231 cells were cultured in DMEM (Cat. No. 10-013-CV, Corning) supplemented with 10% Fetal Bovine Serum (FBS, Cat. No. 97068-085, Avantor) and 1% Penicillin-Streptomycin-Glutamine (PSG, Cat. No. 10378-016, Gibco). PY8119 cells were cultured in Ham’s F-12K Medium (Cat. No. 21127-022, Sigma) with 5% FBS and 1% PSG. PyMT-2900 cells were cultured in a 1:1 mix of DMEM and Ham’s F-12 Nutrient Mix (Cat. No. 11765-054, Gibco) supplemented with 2% FBS, 1% Bovine Serum Albumin (Cat. No. A3803, Sigma), 1% PSG, 10 ng/ml Human EGF Recombinant Protein (Cat. No. PHG0311, Gibco), and 10 μg/ml Insulin (Cat. No. 12585014, Gibco). All cell lines were used between passage 5 and passage 18 and routinely checked for mycoplasma with Universal Mycoplasma Detection Kit (Cat. No. 301012K, ATCC). All cell lines used in this study were mycoplasma negative.

### Conditioned media

Conditioned media was collected from MCF10A, MDA-MB-231, and PY8119 cells for neuron morphology experiments. Briefly, cells were grown to 80% confluency in their respective cell culture medias (as described above). Cells were split 1:5 and cultured in complete neurobasal media. After 48 hours, media was collected from plates, centrifuged at 1500 rpm for 3 min to remove debris and dead cells, and the supernatant was passed through a 0.45 μm Millex MCE Syringe Filter (Cat. No. SLHAR33SS, Millipore).

### Co-culture experiments

DRG dissections were performed on 6- to 10-week-old female C57BL/6J (Strain #000664, Jackson Laboratory), *Trpv1^-/-^*(Strain #003770, Jackson Laboratory), or *Il6^-/-^* (Strain #002650, Jackson Laboratory) mice. DRG were dissociated, resuspended in complete neurobasal media, and seeded at 5,000 – 7,000 cells per well in 24-well plates coated with PDL and laminin, as described above. 24 hours after seeding, a full media change was performed with complete neurobasal media. 48 to 72 hours after seeding, media was fully replaced with either 100% complete neurobasal media (DRG only), a 1:1 mixture of conditioned media and complete neurobasal media (DRG + CM), or 100% complete neurobasal media containing resuspended cells (DRG + cells). A half media change was performed every 48 hours for the duration of the experiment. For conditioned media wells, each half media change consisted of a 1:1 mixture of conditioned media and complete neurobasal media. For experiments involving PY8119 cells, all conditions were maintained in 25% PY8119 culture media (Ham’s F-12K supplemented with 5% FBS and 1% PSG) , as PY8119 cells did not survive in 100% complete neurobasal media.

### 2D immunofluorescence staining and analysis

For 2D experiments, 24-well plates were fixed in 4% Paraformaldehyde (Cat. No. 15710-S, Electron Microscopy Sciences) in PBS for 20 min then permeabilized with 0.1% TritonX-100 (Cat. No. T9284, Sigma), blocked with 6% Normal Donkey Serum (Cat. No. 017-000-121, Jackson ImmunoResearch), and incubated with primary antibodies (Table 1) overnight at 4°C. The following day, wells were incubated with DAPI (dilution: 1/1000, Cat. No. D1306, Invitrogen) and fluorophore-conjugated donkey secondary antibodies (dilution: 1/500, Jackson ImmunoResearch), including Alexa Flour 488, Alexa Flour 545, and Alexa Flour 647, overnight at 4°C. Imaging was performed using a Keyence BZ-X710 microscope at 20X or 40X magnification. For immunofluorescence staining, CellProfiler v4.2.5 was used to quantify nuclear fluorescence intensity (ATF3, cFos, cJun) in DRG neurons using a custom pipeline, with β3-tubulin identifying neurons and DAPI defining nuclear masks, and ImageJ was used to quantify fluorescence intensity (CGRP, TRPV1) in DRG neuron cell bodies. For morphological analysis of neurons, images containing a single neuron cell body per field of view were acquired and analyzed using ImageJ. Various morphological characteristics were assessed, including number of primary neurites, neurite maximum diameter, neuron cell body area, neurite bundling, and bundle maximum diameter. Primary neurites were defined as neuronal processes emanating from a neuron cell body with a length at least twice the diameter of the corresponding neuron cell body. Neurite bundling was defined as two or more processes traveling in parallel for greater than 60 μm. Bundling was quantified as the percentage of primary neurites that exhibit bundling, with each data point representing a single biological replicate. Each analysis was performed at a defined timepoint following initiation of co-culture: cJun at day 4, cFos at day 8, ATF3, CGRP, TRPV1, and morphological assessments at day 12.

### Microelectrode arrays

Microelectrode array (MEA) recordings were performed using the MaxOne High-Density Microelectrode Array (HD-MEA) System (Maxwell Biosystems) to measure DRG neuron electrical activity across conditions. MEA chips were cleaned with 1% Tergazyme solution (Cat. No. 1304-1, Alconox) for 2 hours at room temperature then washed three times with deionized water. Chips were then incubated in 70% ethanol for 30 minutes at room temperature and washed six times with deionized water. Complete neurobasal media was added to the chips for 48 hours to allow for pre-conditioning. 24 hours prior to plating, 50 μl of 0.5 mg/ml Poly-D-Lysine (Cat. No. P0899, Sigma) in deionized water was applied to the electrode region of the chip. The next day, DRG neurons were isolated and dissociated from 6-week-old female SW-F (Taconic) or *Trpv1^-/-^* (Strain #003770, Jackson Laboratory), following the same procedures described above. A suspension of 2,200 cells/μl in complete neurobasal media was mixed with Laminin (Cat. No. L2020, Sigma) for a final plating mixture of 99,000 cells and 0.02 mg/ml Laminin in 50 μL, which was pipetted directly onto the electrode field. One hour after plating, complete neurobasal media was gently added to allow neuronal adherence. After 48 hours, 2,000 MCF10A, MDA-MB-231, or PY8119 cells were added per chip for co-culture experiments. Half media changes were performed every 48 hours, and a final half media change was completed 4 hours prior to recording. Recordings were conducted 21 days post-plating using a gain of 512x, high-pass filter at 300 Hz, and a spike detection threshold of 5.0 for 5 minutes. Data was analyzed using a custom MATLAB script focused on spiking activity per electrode.

### 3D hydrogels

DRG were dissected from 6- to 10-week-old female C57BL/6J (Strain #000664, Jackson Laboratory) or *Trpv1^-/-^*(Strain #003770, Jackson Laboratory) mice and placed on ice in Hibernate-A Medium (Cat. No. A1247501, Gibco). Hydrogels were fabricated as previous described, with modifications^87^. Briefly, hydrogels were composed of 1.7 mg/mL Collagen Type I, Rat Tail (Cat. No. 08-115, Sigma), 10 μg/mL Hyaluronic Acid (Cat. No. 385908, Millipore), 20 μg/mL Laminin Mouse Protein (Cat. No. 23017-015, Sigma), and 20 μg/mL Human Fibronectin (Cat. No. F1056, Sigma), adjusted to pH 7.3 with 0.1 N NaOH (Cat. No. S2770, Sigma). 200 μL of solution was plated in each well of a four-chamber slide (Cat. No. 354104, Corning), and a DRG explant was placed in the middle using a p10 pipette tip. Hydrogels were incubated at 37°C for 1 hour to polymerize before adding 1 mL of complete neurobasal media per well and gently detaching gels. A half media change was performed every 48 hours for the duration of the experiment.

For immunofluorescence staining, hydrogels were washed with PBS, fixed for 30 min at room temperature in 4% Paraformaldehyde (Cat. No. 15710-S, Electron Microscopy Sciences) in PBS, and washed in 0.05% Tween-20 (Cat. No. P1379, Sigma) in PBS (PBS-T). Samples were then permeabilized overnight at room temperature in 0.1% Triton X-100 (Cat. No. T9284, Sigma) in PBS-T, blocked for 2 hours at room temperature in 3% Bovine Serum Albumin (Cat. No. A3803, Sigma) and 3% Normal Donkey Serum (Cat. No. 017-000-121, Jackson ImmunoResearch) in PBS, and incubated with Anti-β3-Tubulin (Table 1) overnight at 4°C. The following day, samples were washed with PBS-T and incubated with Alexa Flour 594 AffiniPure Donkey Anti-Rabbit IgG (Cat. No. 711-585-152, Jackson ImmunoResearch) for 2 hours followed by 5 μg/mL DAPI (Cat. No. D1306, Invitrogen). Hydrogels were then mounted on microscopy slides with ProLong Diamond Antifade Mountant (Cat. No. P36970, Invitrogen) and partially flattened with a coverslip. Imaging was performed using a Zeiss LSM900 confocal microscope (Zeiss) at 5X magnification with stitching to capture the entire hydrogel.

For directionality experiments, 5 μL of hydrogel solution containing 400 MDA-MB-231 cells was plated on one end of a well in a four-chamber slide, while another 5 μL of hydrogel solution was plated in the middle of the well, in which a DRG explant was placed inside. Hydrogels were incubated at 37°C for 10-15 min to begin solidification, after which 200 uL of hydrogel solution was carefully placed on top. After another 1 hour of incubation at 37°C, 1 mL of complete neurobasal media was added per well. Hydrogels were cultured for 16-20 days. Media changes and immunofluorescence staining were completed as described above.

### RNA sequencing and analysis

RNA-seq data analyzed in this study were previously generated by the Oudin lab to investigate the effects of DRG on breast cancer cells and made publicly available in GEO under accession code GSE180508^15^. Briefly, DRG neurons were cultured alone or co-cultured with MDA-MB-231 cells for 24 hours, after which total RNA was extracted using the RNA MiniPrep Kit (Cat. No. R1057, Zymo Research). Libraries were prepared using the TruSeq Stranded mRNA kit and sequenced by the Tufts Genomics Core on the NextSeq550 platform at read length of 150 nt, paired-end, with sequencing depths ranging from 28 to 35 million reads per mono-culture samples and 55 to 72 million reads per co-culture samples. Reads were subjected to quality control using FastQC^88^, followed by adapter trimming and filtering with Trimmomatic^89^. Reads were aligned to both the human (GRCh38) and mouse (GRCm38) genomes using STAR^90^, and species-specific transcriptomes from co-culture samples were deconvolved using the S3 algorithm^91^. DRG mono-culture samples were processed in parallel using the same pipeline to ensure consistency. Gene expression analysis was performed using the EdgeR^92^ package in R. Genes with counts per million (CPM) <1 were excluded from analysis. Principal component analysis (PCA) was performed using the prcomp() function in R on the TPM values of filtered genes. Differentially expressed genes (DEGs) between DRG mono-culture and DRG co-cultured with MDA-MB-231 were identified using a fold-change cutoff of 1.2. Pathway enrichment analysis on DEGs between DRG and DRG co-cultured with MDA-MB-231 was performed using GSEA^93^ and GOrilla^94^ to identify enriched Gene Ontology (GO) biological processes and pathways and summarized using Revigo^95^. Transcription factor enrichment analysis was performed using the TRRUST^96^ function on Metascape (metascape.com, RRID: SCR_016620) based on differentially expressed genes with a fold-change cutoff of 1.2. The top 15 enriched transcription factors were graphed.

### Animal experiment

1×10^6^ PY8119 cells were suspended in 100 μl solution of 20% Collagen I (Cat. No. 354236, Corning) in PBS and injected into the fourth right mammary fat pads of 7-week-old female C57BL/6J (Strain #000664, Jackson Laboratory) or *Trpv1^-/-^*(Strain #003770, Jackson Laboratory) mice. Tumor volume was measured every 3 days by calipers in duplicate starting at day 10, once tumors were palpable, following the formula: tumor volume = [length x width x width]/2. Mice were euthanized by CO_2_ 25-days after injection and tumors and lungs were excised, washed in PBS, and fixed for 24 hours in 4% Paraformaldehyde (Cat. No. 15710-S, Electron Microscopy Sciences) in PBS. Tissues were embedded in paraffin and sectioned into 10 μm sections. For hematoxylin and eosin staining of lung tissue, sections were deparaffinized, hydrated, and stained using standard procedures with Hematoxylin (Cat. No. GHS280, Sigma) and counterstained with Eosin (Cat. No. HT110180, Sigma). Lung metastases were quantified by two observers.

### Statistical analysis

Results were analyzed with GraphPad Prism software (version 10.1.1) using unpaired Student’s t tests, one-way ANOVA, or Fisher’s exact tests. Results are displayed as mean ± SEM. *p<0.05, **p<0.01, ***p<0.001, ****p<0.0001.

## Author Contributions

H.B., S.R.P., M.L., C.K., and M.J.O designed the experiments. S.R.P. performed retrograde labeling and S.R.P. and H.B. analyzed results. H.B., M.M.C., and T.A. performed in vitro staining and fluorescence intensity analysis. H.B. performed in vitro staining and morphological analysis. A.L.P. and H.B. performed DRG histology. T.G. wrote script to assist in DRG histology analysis and A.L.P. performed analysis. H.B. and A.P. performed DRG qPCR. S.R.P. and W.C. performed and analyzed microelectrode array experiments. H.B. and T.T.L. performed RNA-seq analysis. H.B. and S.R.P. performed and analyzed in vivo experiment. H.B., S.R.P., and M.J.O. wrote and edited the manuscript.

## Acknowledgements

H.B. acknowledges support by NIH Grant No. 1F30 CA294624-01 and NIH Grant No. 5T32 GM008448-27. M.J.O. acknowledges support by NIH Grant No. R01CA255742 and DP2CA271387 and the Breast Cancer Alliance. W.C. acknowledges support by AFOSR Grant No. FA9550-22-1-0465. The authors thank all members of the Oudin Lab for helpful discussions; Dr. Colin Trepicchio, Meadow Parrish, and Nicole Traugh from the Kuperwasser Lab and Dr. Daniel Miller from Piyush Gupta’s Lab for help with 3D hydrogel cultures; the Tufts Comparative Medicine Services personnel for animal care; the Tufts Comparative Medicine Services Histology Core for tissue processing; and the Tufts Flow Cytometry Core for help in generating GFP+ cell lines.

Supplemental Materials | Bloomer et al.

## Supplemental Materials

**Figure S1.**
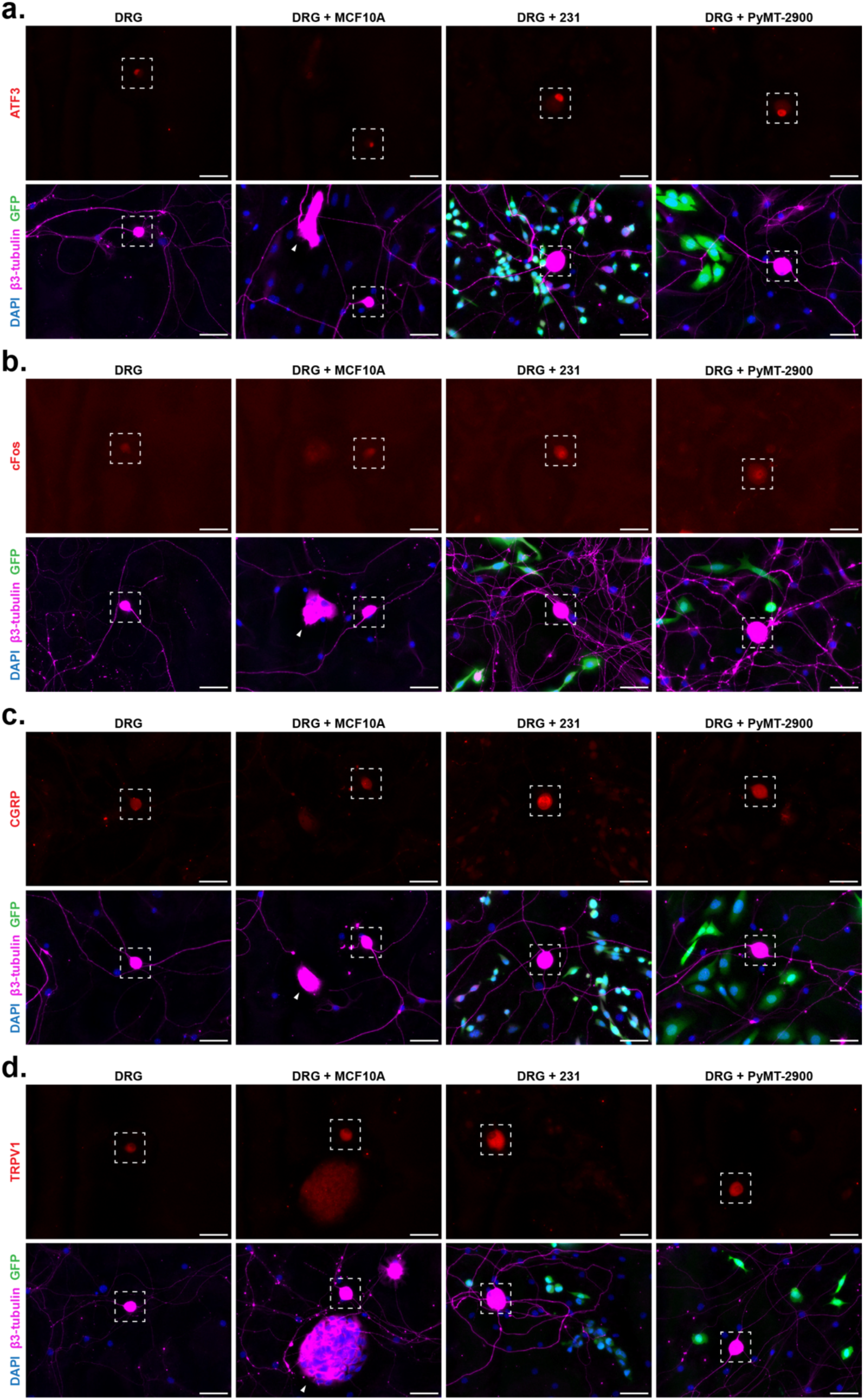
Representative images of DRG from C57BL/6J mice cultured alone or co-cultured with MCF10A cells, GFP+ MDA-MB-231 (231) cells, or GFP+ PyMT-2900 cells. Cells were stained for DAPI, β3-tubulin, and (**a**) ATF3, (**b**) cFos, (**c**) CGRP, and (**d**) TRPV1. White dashed box corresponds to DRG neuron cell body and representative images in Figure 1E, 1G, 1I and 2A. White arrow indicates MCF10A cells. Scale, 50 µm.

**Figure S2.**
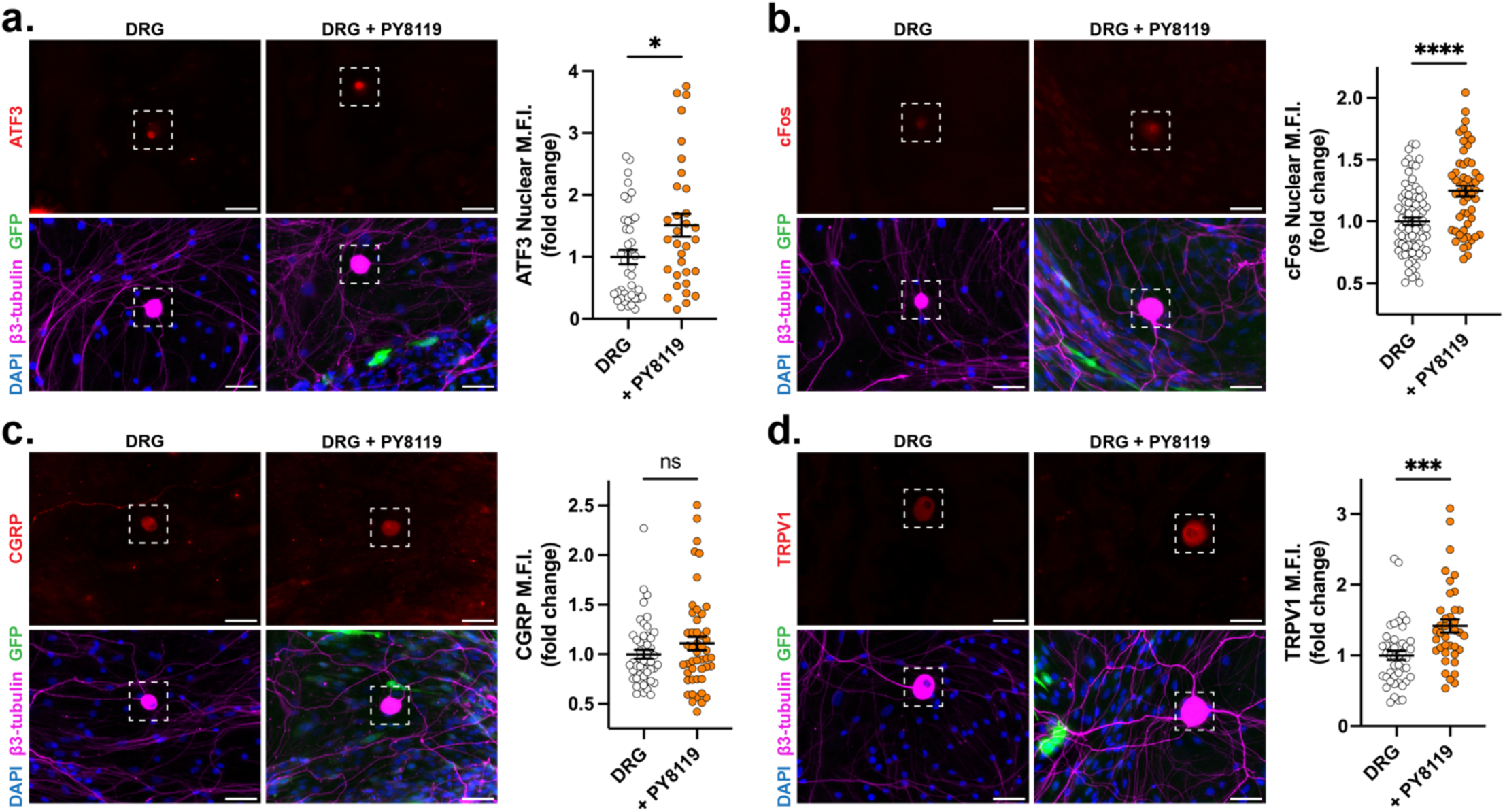
Representative images and analysis of DRG from C57BL/6J mice cultured alone or co-cultured with GFP+ PY8119 cells. Cells were stained for DAPI, β3-tubulin, and various activation markers. **a)** Nuclear ATF3 mean fluorescence intensity (MFI) in DRG cultured alone (n=3 biological replicates, 41 neurons) or co-cultured with GFP+ murine PY8119 cells (n=3, 32 neurons) for 12 days. **b)** Nuclear cFos MFI in DRG cultured alone (n=4, 81 neurons) or co-cultured with GFP+ murine PY8119 cells (n=4, 54 neurons) for 8 days. **c)** CGRP MFI in DRG neuron cell bodies when cultured alone (n=3, 49 neurons) or co-cultured with GFP+ murine PY8119 cells (n=3, 48 neurons) for 12 days. **d)** TRPV1 MFI in DRG neuron cell bodies when cultured alone (n=3, 45 neurons) or co-cultured with GFP+ murine PY8119 cells (n=3, 38 neurons) for 12 days. White dashed box corresponds to DRG neuron cell body. Scale, 50 µm. Data presented as mean ± SEM. Statistical significance determined by Student’s t-test, *p<0.05, ***p<0.001, ****p<0.0001.

**Figure S3.**
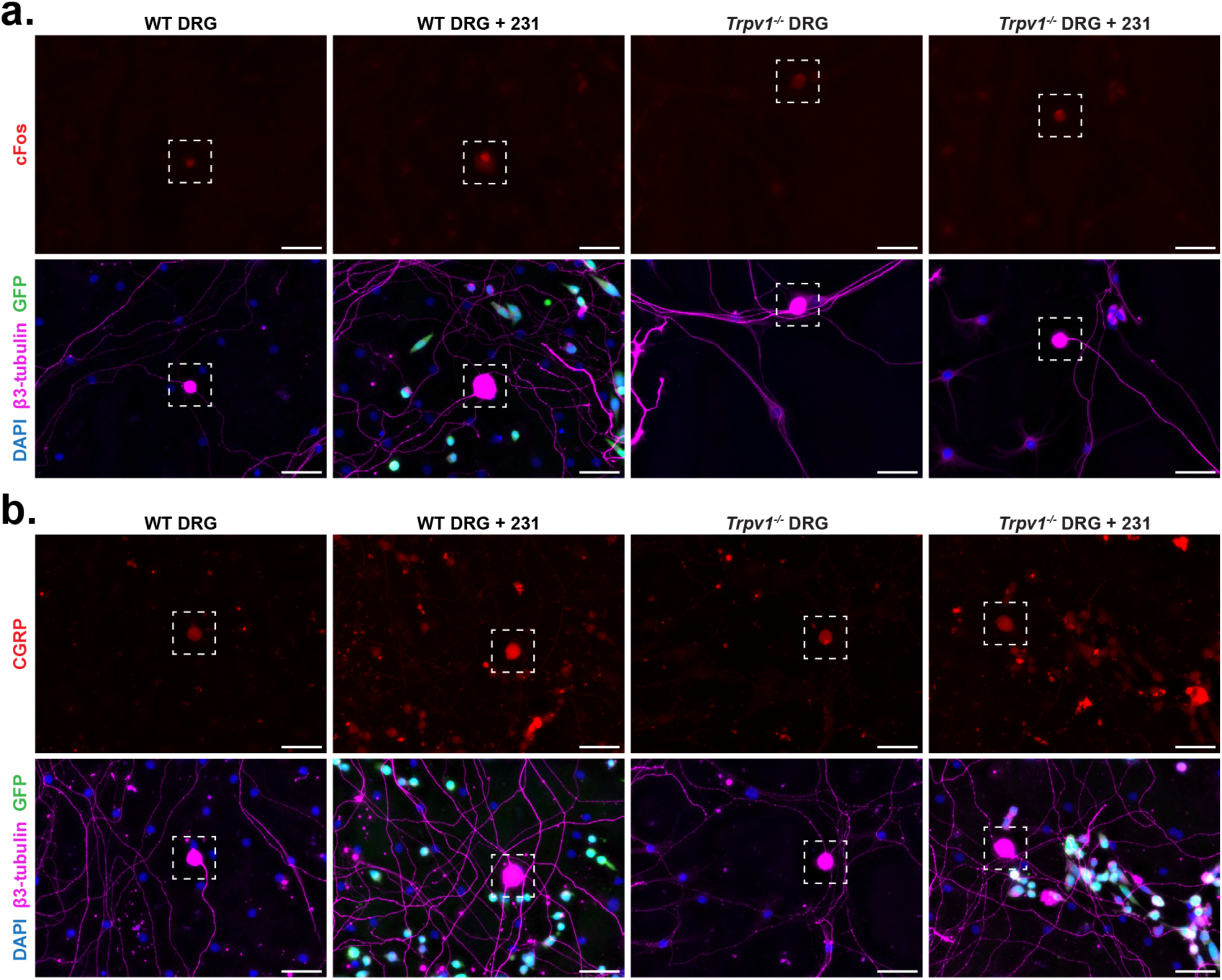
Representative images of DRG from C57BL/6J (WT) or *Trpv1^-/-^* mice cultured alone or co-cultured with GFP+ MDA-MB-231 (231) cells. Cells were stained for DAPI, β3-tubulin, and (**a**) cFos or (**b**) CGRP. White dashed box corresponds to DRG neuron cell body and representative images in Figure 2B and 2C. Scale, 50 µm.

**Figure S4.**
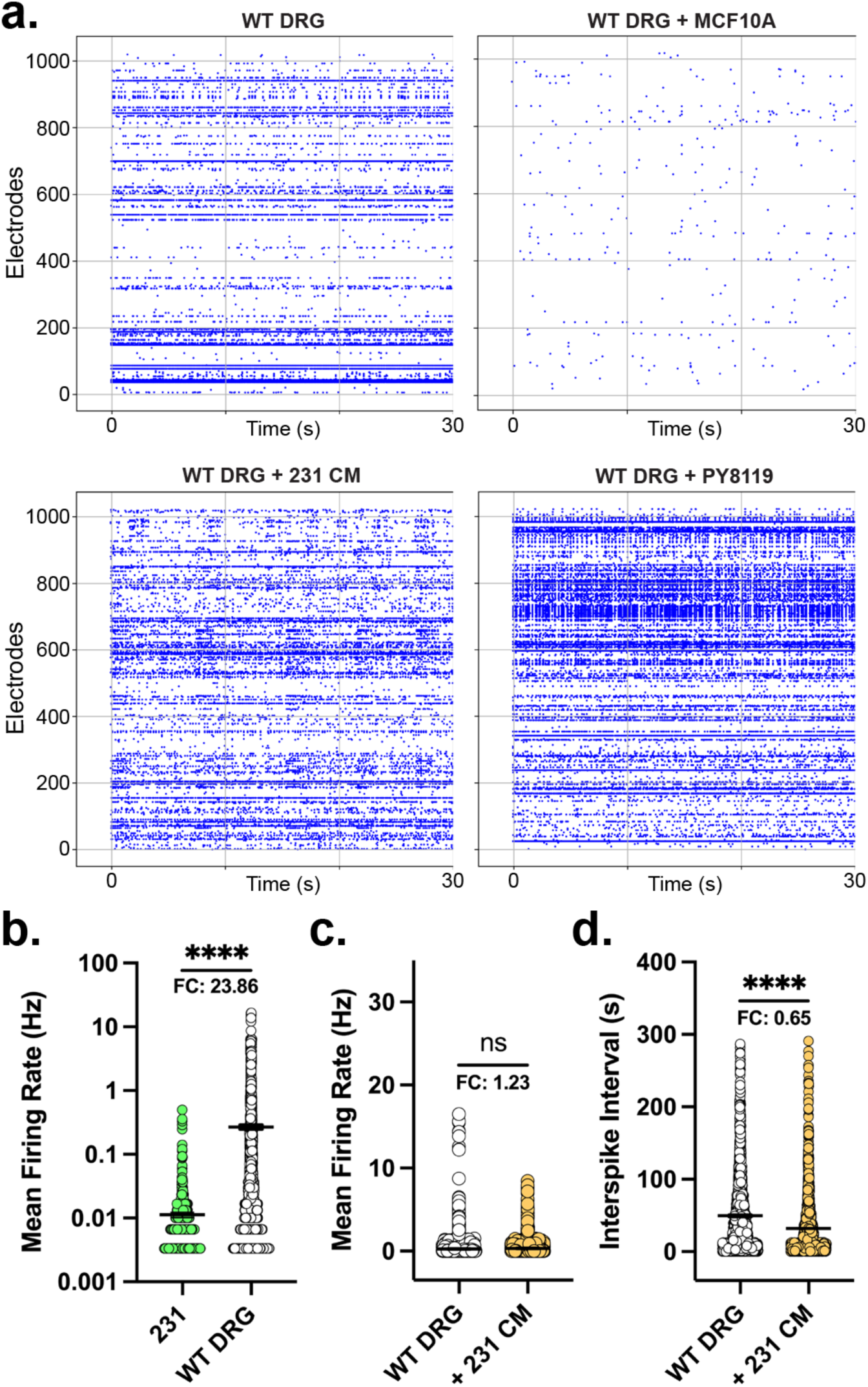
Cells were cultured for 21 days on microelectrode arrays (MEAs) and bioelectric activity was measured for 300 seconds at endpoint. **a)** Representative raster plots of DRG from SF-W (WT) mice cultured alone, with conditioned media (CM) from MDA-MB-231 (231) cells, or co-cultured with MCF10A or PY8119 cells. Each data point represents a detectable spike within the 30 second window. **b)** Mean firing rate, the number of spikes per second per electrode (Hz), in 231 cells cultured alone (n=1) or WT DRG cultured alone (n=4). **c)** Mean firing rate (Hz) in WT DRG cultured alone (n=4) or cultured with CM from 231 cells (n=2). **d)** Interspike interval is plotted as the average time between two spikes per electrode. Data presented as mean ± SEM. Statistical significance determined by Student’s t-test, ****p<0.0001.

**Figure S5.**
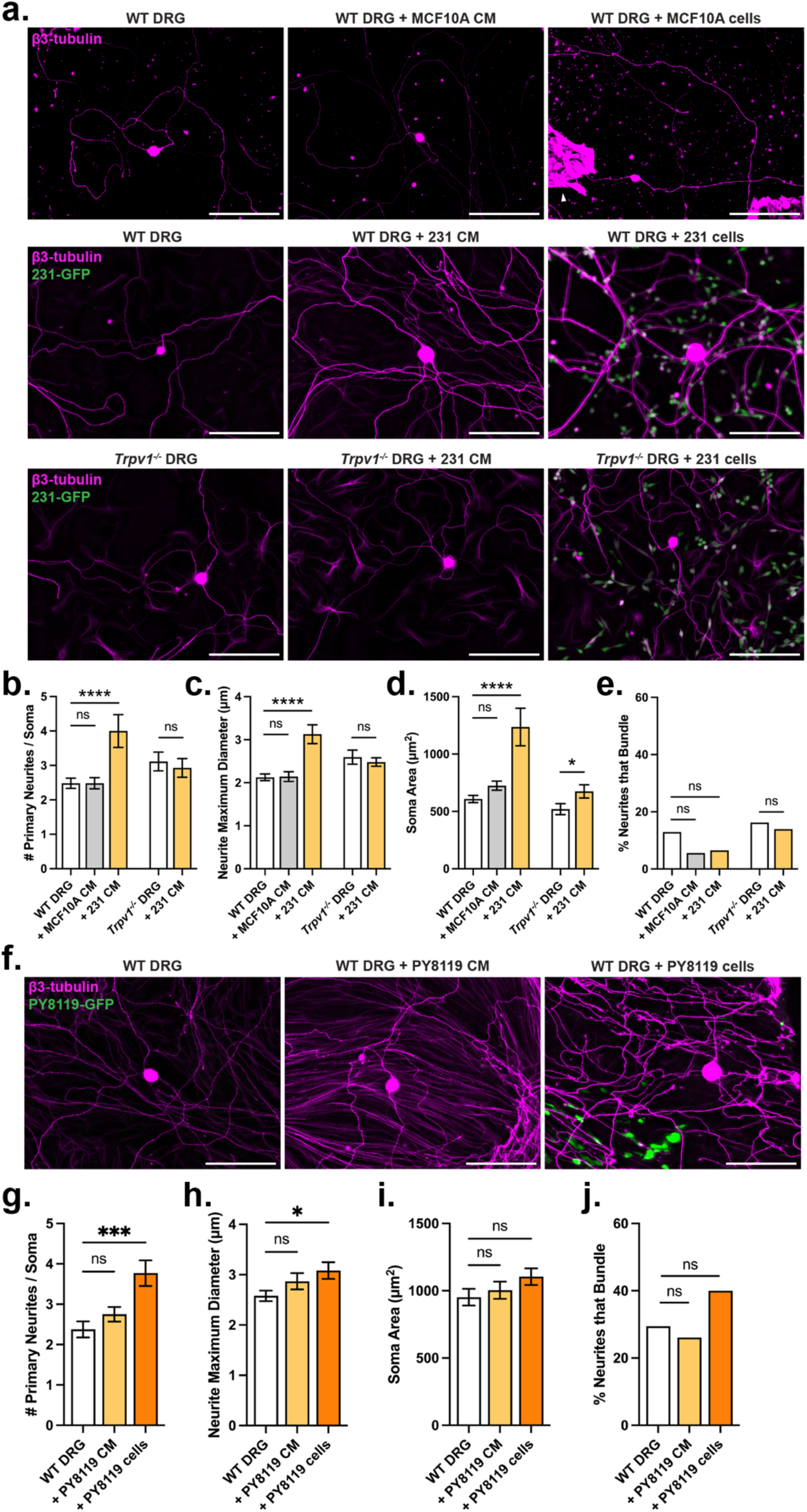
**a)** Representative images of C57BL/6J (WT) or *Trpv1^-/-^* DRG cultured alone, with conditioned media (CM) from MCF10A or MDA-MB-231 (231) cells, or co-cultured with MCF10A or GFP+ 231 cells and stained for β3-tubulin. White arrow indicates MCF10A cells. Scale bar, 200 μm. **b-e)** WT or *Trpv1^-/-^* DRG cultured alone or with conditioned media from MCF10A or 231 cells were analyzed for various morphological features linked to neurite initiation and growth by ImageJ. **b)** Number of primary neurites per soma. WT DRG cultured alone (n=8 biological replicates, 56 neurons) or with MCF10A CM (n=4, 29 neurons) or 231 CM (n=2, 16 neurons). *Trpv1^-/-^*DRG cultured alone (n=4, 26 neurons) or with 231 CM (n=4, 28 neurons). **c)** Neurite maximum diameter per field of view (FOV). WT DRG cultured alone (n=6, 39 FOV) or with MCF10A CM (n=2, 16 FOV) or 231 CM (n=2, 12 FOV). *Trpv1^-/-^* DRG cultured alone (n=4, 26 FOV) or with 231 CM (n=4, 27 FOV). **d)** Soma area. WT DRG cultured alone (n=8, 56 neurons) or with MCF10A CM (n=4, 29 neurons) or 231 CM (n=2, 16 neurons). *Trpv1^-/-^*DRG cultured alone (n=4, 26 neurons) or with 231 CM (n=4, 28 neurons). **e)** Percent of neurites that bundle. WT DRG cultured alone (n=8, 18/139) or with MCF10A CM (n=4, 4/72) or 231 CM (n=2, 4/61). *Trpv1^-/-^* DRG cultured alone (n=4, 13/80) or with 231 CM (n=4, 11/79). **f)** Representative images of WT DRG cultured alone, with conditioned media (CM) from PY8119 cells, or co-cultured with GFP+ PY8119 cells and stained for β3-tubulin. Scale bar, 200 μm. **g-j)** DRG were analyzed for various morphological features linked to neurite initiation and growth by ImageJ. **g)** Number of primary neurites per soma. WT DRG cultured alone (n=40, 28 neurons), with PY8119 CM (n=4, 40 neurons), or co-cultured with PY8119 cells (n=4, 39 neurons). **h)** Neurite maximum diameter per field of view (FOV). WT DRG cultured alone (n=2, 20 FOV), with PY8119 CM (n=2, 20 FOV), or co-cultured with PY8119 cells (n=2, 20 FOV). **i)** Soma area. Number of primary neurites per soma. WT DRG cultured alone (n=4, 40 neurons), with PY8119 CM (n=4, 40 neurons), or co-cultured with PY8119 cells (n=4, 39 neurons). **j)** Percent of neurites that bundle. WT DRG cultured alone (n=4, 28/95), with PY8119 CM (n=4, 29/111), or co-cultured with PY8119 cells (n=4, 58/145). Data presented as mean ± SEM. Statistical significance determined by Student’s t-test (b-d), one-way ANOVA (b-d, g-i) or Fisher’s exact test (e, j), *p<0.05, **p<0.01, ***p<0.001, ****p<0.0001.

**Figure S6.**
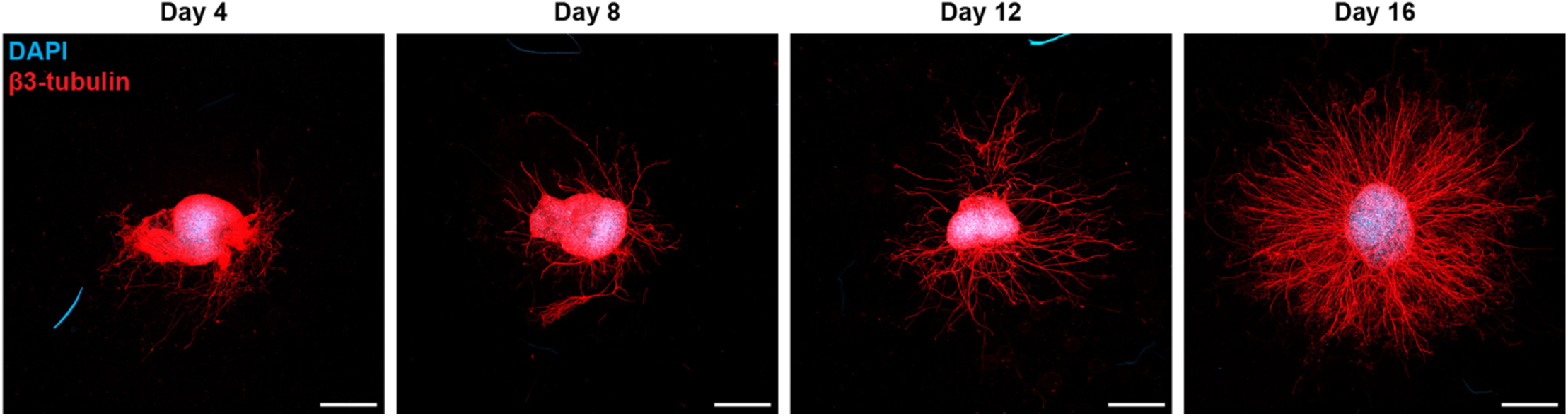
DRG explants from C57BL/6J mice were cultured in 3D hydrogels for up to 16 days and stained for DAPI and β3-tubulin. Scale, 500 µm.

**Figure S7.**
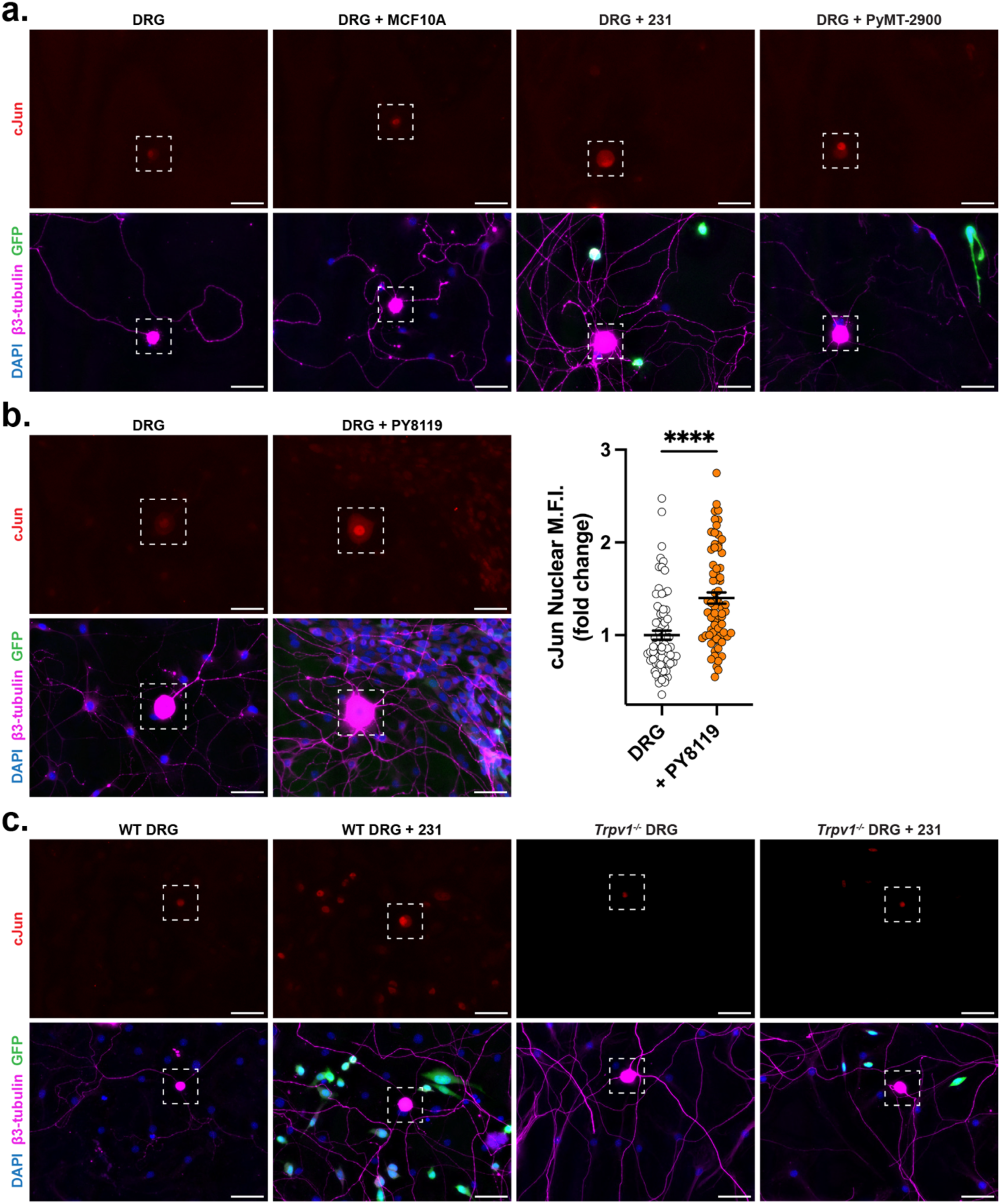
**a)** Representative images of DRG from C57BL/6J mice cultured alone or co-cultured with MCF10A cells, GFP+ MDA-MB-231 (231) cells, or GFP+ PyMT-2900 cells. Cells were stained for DAPI, β3-tubulin, and cJun. White dashed box corresponds to DRG neuron cell body and representative images in Figure 5E. Scale, 50 µm. **b)** Representative images and analysis of DRG from C57BL/6J mice cultured alone or co-cultured with GFP+ PY8119 cells. Cells were stained for DAPI, β3-tubulin, and cJun. Nuclear cJun mean fluorescence intensity (MFI) in DRG cultured alone (n=6 biological replicates, 75 neurons) or co-cultured with GFP+ murine PY8119 cells (n=6, 71 neurons) for 4 days. White dashed box corresponds to DRG neuron cell body. Scale, 50 µm. Data presented as mean ± SEM. Statistical significance determined by Student’s t-test, ****p<0.0001. **c)** Representative images of DRG from C57BL/6J (WT) or *Trpv1^-/-^* mice cultured alone or co-cultured with GFP+ MDA-MB-231 (231) cells. Cells were stained for DAPI, β3-tubulin, and cJun. White dashed box corresponds to DRG neuron cell body and representative images in Figure 5F. Scale, 50 µm.

**Figure S8.**
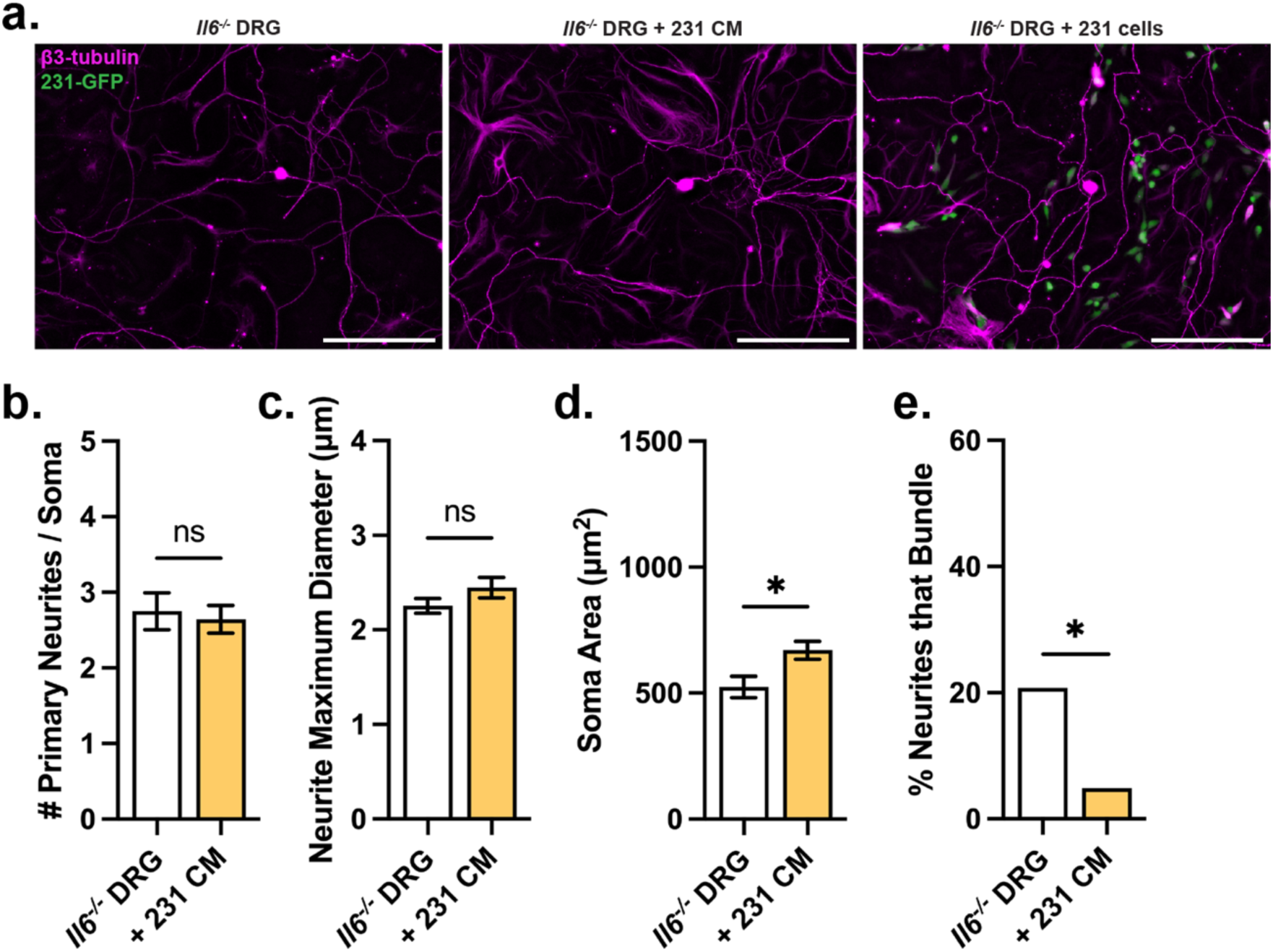
**a)** Representative images of *Il6^-/-^* DRG cultured alone, with conditioned media (CM) from MDA-MB-231 (231) cells, or co-cultured with GFP+ 231 cells and stained for β3-tubulin. Scale bar, 200 μm. **b-e)** DRG were analyzed for various morphological features linked to neurite initiation and growth by ImageJ. **b)** Number of primary neurites per soma. *Il6^-/-^* DRG cultured alone (n=4 biological replicates, 28 neurons) or with 231 CM (n=4, 31 neurons). **c)** Neurite maximum diameter per field of view (FOV). *Il6^-/-^*DRG cultured alone (n=2, 16 FOV) or with 231 CM (n=2, 16 FOV). **d)** Soma area. Number of primary neurites per soma. *Il6^-/-^* DRG cultured alone (n=4, 28 neurons) or with 231 CM (n=4, 32 neurons). **e)** Percent of neurites that bundle. *Il6^-/-^* DRG cultured alone (n=4, 16/77) or with 231 CM (n=4, 4/82 neurons). Data presented as mean ± SEM. Statistical significance determined by Student’s t-test (b-d) or Fisher’s exact test (e), *p<0.05.

